# Predicting missing links in food webs using stacked models and species traits

**DOI:** 10.1101/2024.11.22.624890

**Authors:** Lucy Van Kleunen, Laura E. Dee, Kate L. Wootton, François Massol, Aaron Clauset

## Abstract

Networks are a powerful way to represent the complexity of complex ecological systems. However, most ecological networks are incompletely observed, e.g., food webs typically contain only partial lists of species interactions. Computational methods for inferring such missing links from observed networks can facilitate field work and investigations of the ecological processes that shape food webs. Here, we describe a stacked generalization approach to predicting missing links in food webs that can learn to optimally combine both structural and trait-based predictions, while accounting for link direction and ecological assumptions. Tests of this method on synthetic food webs show that it performs very well on networks with strong group structure, strong trait structure, and various combinations thereof. Applied to a global database of 290 food webs, the method often achieves near-perfect performance for missing link prediction, and performs better when it can exploit both species traits and patterns in connectivity. Furthermore, we find that link predictability varies with ecosystem type, correlates with certain network characteristics like size, and is principally driven by a subset of ecologically-interpretable predictors. These results indicate broad applicability of stacked generalization for studying ecological interactions and understanding the processes that drive link formation in food webs.

## I. INTRODUCTION

Many complex social, technological, and biological systems can be represented as networks, defined as a set of nodes, e.g., individual species, people, genes, or even places, along with their pairwise interactions, e.g., feeding relationships between species in food webs, friendships in social networks, regulatory interactions between genes, or traffic flows among places. However, nearly all empirical networks are incomplete, because real links can be unobserved, unmeasured, hidden, or inaccessible. For example, in food webs representing species feeding relationships, both hard to observe feeding events and rare interactions may be missing. Although many methods now exist for inferring such “missing links” based on their correlation with a network’s partially observed structure [13, 28, 39, 44, 66, 68, 69, 115], we lack highly-accurate methods that leverage the particular characteristics of food webs to make ecologically accurate predictions of species interactions.

Broadly, link prediction methods based on network structure can be grouped into three classes [39]: those that predict missing links based on (i) the pattern of links local to where a missing link may occur, (ii) large-scale models of the entire network’s structure (e.g., grouping structure), and (iii) node proximity within a learned embedding of the network. Systematic evaluations of link prediction methods using large corpora of structurally diverse empirical networks indicate that there is no universally best method for all networks [39], and the best approach depends on the particular network. Among modern link prediction methods, the meta-learning approach of stacked generalization [112], or model stacking, is a state-of-the-art technique that can learn from patterns among observed network interactions how to optimally combine many individual link predictors to produce highly accurate predictions in real-world social, biological, and technological networks [39]. Using existing stacking methods, missing links in social networks are the easiest to recover, while missing links in biological networks, including food webs, remain substantially harder to predict [39].

Hence, tailored approaches are required in particular domains like predicting missing links in food webs. Food webs are often used as models of ecosystem structure to assess ecosystem vulnerability to disturbances in theoretical studies and applied conservation contexts [33, 47, 58, 70]. Assembling a food web is labor-intensive, often requiring researchers to carefully identify and combine feeding interactions recorded in the literature with new field observations or experimental results [55, 108], as well as collect or assemble trait data for member species. Feeding links between species can be identified via a number of methods, including expert elicitation, direct observation in the field or in the lab, and molecular analyses of gut content, feces, tissues, or museum specimens. However, because the number of possible feeding links grows quadratically with the number of species considered, while the number of true feeding links typically grows only linearly, distinguishing every true link from all true non-links in even a modest-sized food web can be prohibitive. Hence, the links present in most food web datasets are incompletely sampled [25, 54, 106]. More accurate methods for predicting missing links in food webs would both increase the efficiency of collecting species interaction data in the field and provide more reliable insights into questions about ecosystem stability, conservation efforts, and tests of ecological theories [77].

Food webs have three distinguishing characteristics that existing stacking methods do not account for in missing link prediction. First, species attributes, or traits, like feeding mode, trophic level, body mass, and metabolic type constrain the set of ecologically feasible feeding links [37, 72]. Several studies match species foraging traits with vulnerability traits that constrain interactions (known as “trait matching”) [3, 8, 23, 31, 37, 40, 62, 73, 85, 90, 96, 114]. Second, feeding links are directional. That is, we must not just predict that an interaction exists between two species, but also the correct direction of the feeding interaction and whether it is reciprocated. Third, while food webs are similar to social networks in often exhibiting skewed degree distributions [32, 34] and compartmentalized grouping or community structure [5], they also have structural properties that are different from social networks, in particular exhibiting fewer triangles and a globally hierarchical structure [28, 111]. State-of-the-art model stacking approaches do not currently exploit node attributes or link directionality, and are not customized to expect global hierarchical structure, which limits their utility for making accurate predictions in food webs. Here, we develop a new stacking model specifically designed to exploit these features to make more accurate predictions of missing feeding links.

Food webs provide an ideal setting for exploring how the stacking model approach can be adapted to a specific class of biological networks where interactions are directional. We build on substantial previous work on applying individual link prediction methods to partially observed food webs [30, 31, 79, 87–89, 98, 110] and other ecological networks [28, 38, 72, 78, 87, 93, 98, 106, 114]. Mirroring work on networks in general [39], work on missing link prediction in food webs has found that there is no universally best predictor for missing links in food webs [30, 88, 89, 93, 98, 106, 114]. Meta-learning exploits the fact that many prediction methods work well in practice but do so using complementary underlying signals. By learning to optimally combine these signals, meta-learning can substantially improve prediction accuracies. Here, we build upon past explorations in ecology that have combined prediction methods via averaging, multiplication, or summation of predictors [11, 84, 106]. Model stacking generalizes these ways of combining methods by algorithmically constructing an optimal predictive distribution from individual predictors for a particular data set [112]. Such a meta-learning approach is an attractive strategy for missing link prediction in food webs because it allows us to be relatively agnostic about the theoretical basis of particular distinct predictors, while also using data to guide their combination into a single prediction algorithm that takes advantage of whatever structural regularities are present. In addition, the resulting model can often be interpreted to yield insights into the underlying processes shaping the network [43].

Here, we develop a stacking model for predicting missing links in food webs that combines predictors based on species traits (node attributes) and predictors based on connectivity patterns (network structure), which we adapt to follow ecological assumptions about directed links in food webs. We first evaluate this method using a class of synthetic networks with known structure, which allows us to systematically vary the degree to which links exist due to node attributes or network structure. We then apply the method to a global database of 290 food webs with species trait annotations [19]. We find that model stacking using species traits and connectivity patterns is highly predictive of missing interactions in food webs, and the best performance is generally achieved by combining both types of information. By assessing how performance varies across ecosystem types and network characteristics, we find that missing links are easiest to predict in terrestrial belowground food webs, and in food webs that are larger, have better taxonomic resolution, are more connected, and are less modular.

Further, our results illustrate some of the ecological insights that can be obtained with a method that flexibly learns highly accurate prediction rules for specific food webs. Across 290 food webs, we find that ecosystem type correlates with the relative performance of attribute vs. structure-based predictors as groups, and with which individual predictors are most important for predicting missing links. At the same time, we identify a subset of predictors that are broadly important across ecosystem types, suggesting common underlying processes that structure these networks. This subset includes a custom ecological preferential attachment predictor, the species log_10_ body mass ratio, and predictors based on low rank approximation and ‘nearest neighbors’. These results demonstrate how a model stacking approach that is adapted specifically to ecological networks can produce both highly accurate missing link predictions in food webs and provide new insights into food web organization for the development and verification of ecological theory.

## II. RESULTS

In missing link prediction, the ‘true’ network *G* = (*V, E*) is defined by a set of nodes *V* and a set of edges or links *E*, but is incompletely observed. The observed network *G*^*′*^ = (*V, E*^*′*^) has the same set of nodes *V* but only the ‘observed’ subset of edges *E*^*′*^ ⊂ *E*. Our stacked generalization model for food webs, adapted from Ref. [39] for directed and attributed networks, is described in detail in the Methods section. Briefly, this supervised-learning approach uses an ensemble of missing link predictors, based on network structure and node attributes, to learn how to score all potentially missing (unobserved) links in a particular observed food web *G*^*′*^ such that higher scoring candidates are more likely to be missing links. Hence, this approach defines a network-specific model that learns from a particular network’s observed edges and unique characteristics.

In contrast to past work [39], our stacked model adapts individual link predictors to food webs by grounding them in ecological assumptions. For example, ‘common neighbors’ predicts that a link is more likely to be missing if it would connect a pair of nodes who have many neighbors in common; we modify this idea so that the direction of the predicted connection aligns with ecological assumptions about trophic hierarchies (Fig. 1), and we define a new set of predictors based on assortativity among node attributes (Fig. 2).

**FIG. 1.**
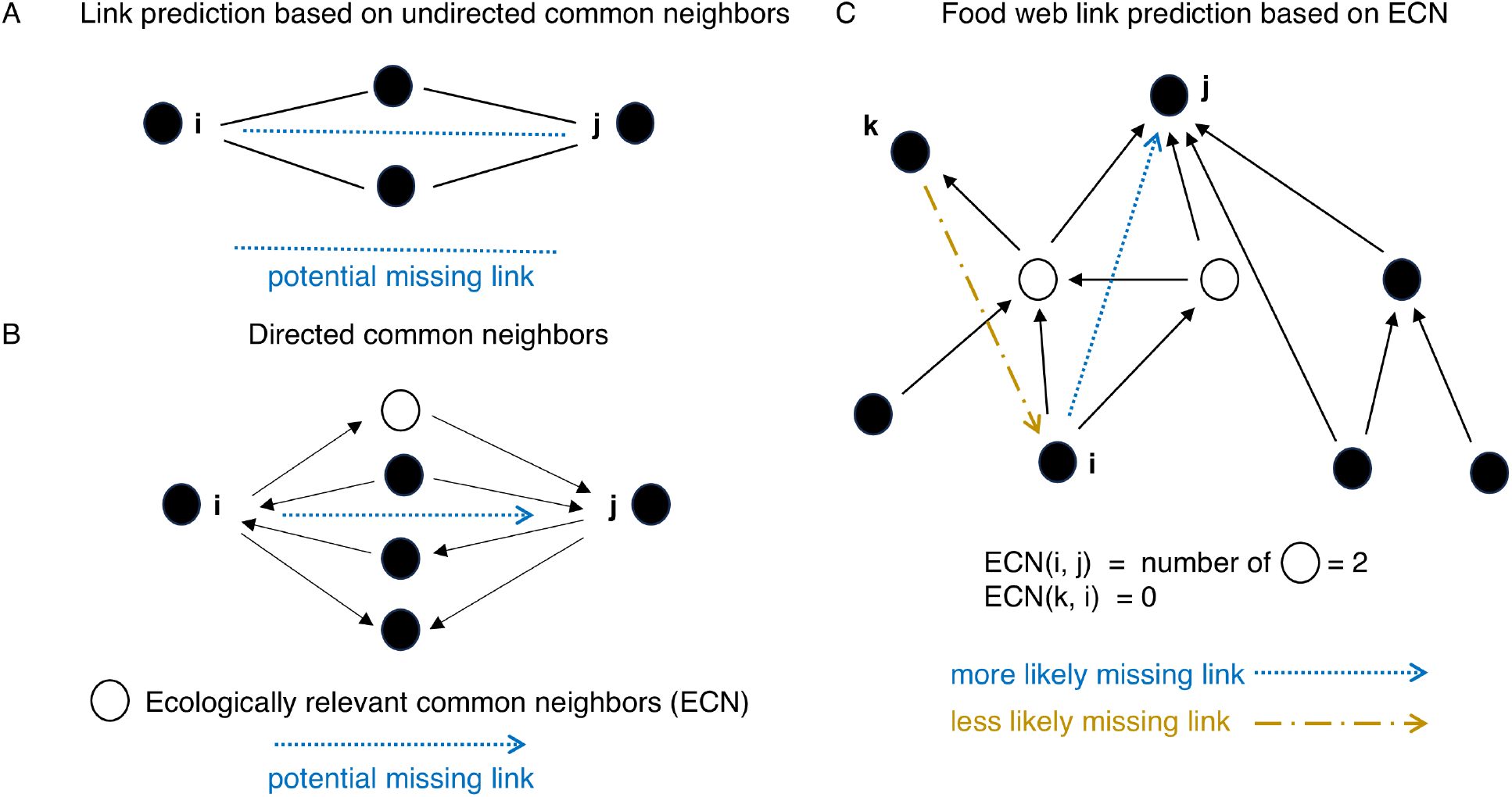
Adapting topological predictors to food webs. (A) In undirected networks, the ‘common neighbors’ predictor assumes that the more neighbors two unconnected nodes *i* and *j* have in common, the greater the likelihood that (*i, j*) is a missing link. (B) For an unconnected pair *i, j* in a directed network, there are four distinct arrangements of directed connections with a common neighbor of *i* and *j*. If there is a missing link between *i* and *j*, the arrangement whose edge directions align with the network’s trophic hierarchy is the more ecologically likely. (C) For example, the ecologically relevant common neighbors (ECN) predictor predicts a missing link (*i, j*) (dashed arrow) because *i, j* have two ecologically relevant common neighbors (open circles), and not the link (*k, i*) (dot-dashed arrow) which have none.

**FIG. 2.**
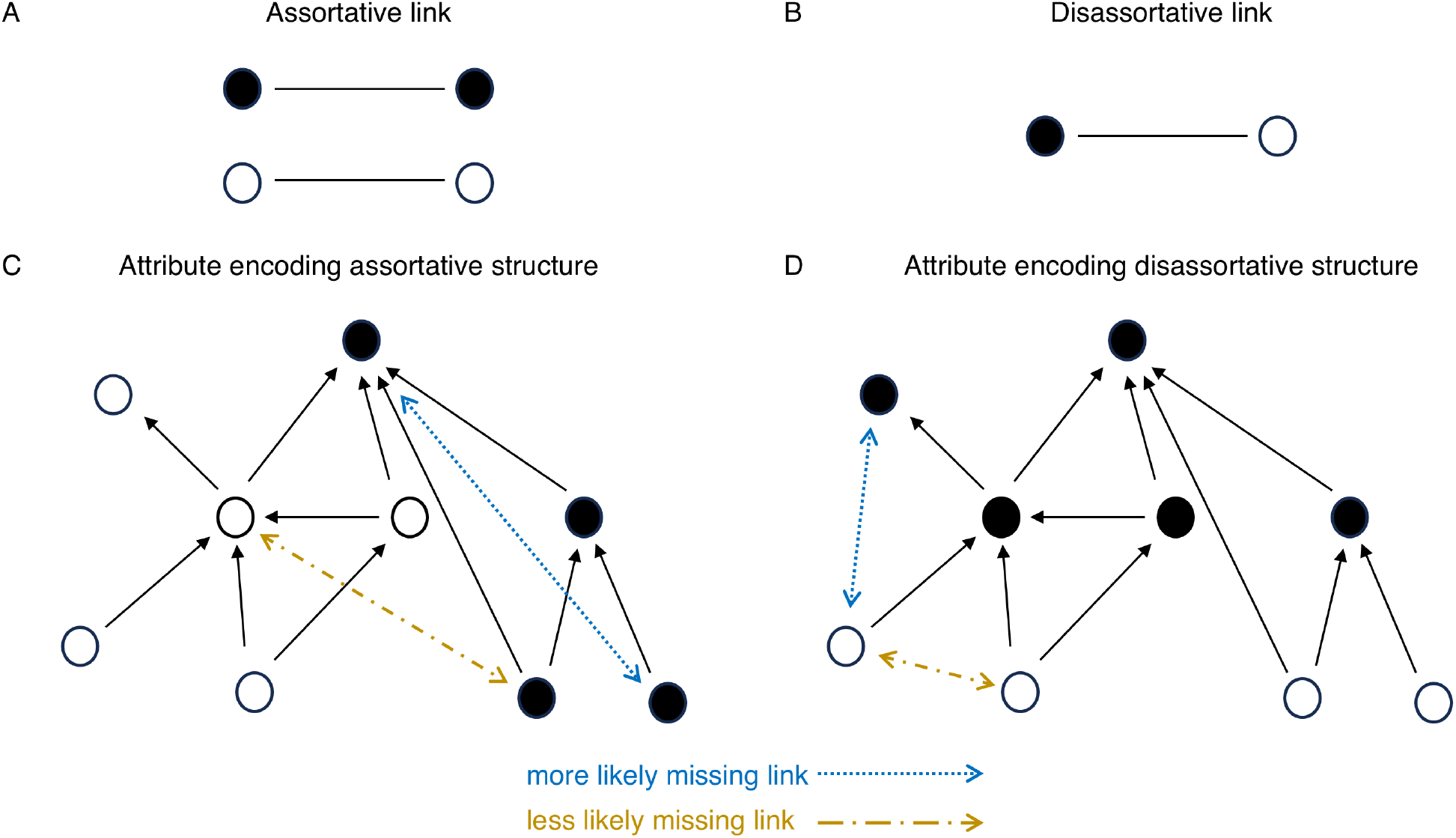
Assortative and disassortative patterns. Node attributes can encode (A) assortative patterns, e.g., common environmental conditions, and (B) disassortative patterns, e.g., a species trait that differs by trophic level, which can be exploited to predict missing links. (C) When node attributes encode assortative information, missing links tend to occur between nodes with similar attributes. (D) In contrast, when node attributes encode disassortative information, missing links will tend to occur between nodes with different attributes. The stacked model (see text) allows us to include both types of attribute predictors and to use data to learn which are most useful in a given network.

### A. Performance on synthetically generated networks

We first evaluate the accuracy of the stacked model using synthetically generated food webs with node attributes. In this controlled setting, the true data generating processes are known, allowing us to calculate the theoretical maximum accuracy, and we can adjust the extent to which the probability of an edge depends on its nodes’ attributes.

In empirical networks with node annotations, observed node attributes often correlate with the network structure, and hence they also correlate with missing links; however, other structural patterns relevant to missing link prediction do not appear to correlate with node attributes [30, 37, 89]. We incorporate these patterns into our synthetic networks by defining a parameter *ρ* ∈ [0, 1], which tunes the probability that an edge’s existence depends on node attributes. When *ρ* = 0, the network structure is completely independent of all node attributes and when *ρ* = 1, the structure is completely determined by node attributes. This range of dependency is accomplished by creating a pair of “anchor” networks with the same number of nodes and approximately the same number of edges, one with strong latent topological structure unrelated to node attributes and one with structure fully determined by node attributes. To generate a network with some mixture of these patterns, *ρ* specifies the fraction of edges sampled from the first anchor network, with the remaining edges sampled from the second.

To generate anchor networks with network structure that is independent of node attributes but with structure that is nevertheless similar to that found in food webs, we use the stochastic block model (SBM) [50]. In the SBM, nodes are assigned to groups and the probability of a link (*i, j*) depends on the group assignments of the nodes *i, j*. To emulate role-based trophic grouping structure in food webs, e.g., between predators, herbivores, and primary producers, the direction of the generated edges were chosen to replicate expected hierarchical structure in food webs (see Materials & Methods). To generate anchor networks with network structure that is fully determined by node attributes, we used the random geometric graph model (RGG) [80] to generate networks with assortative or disassortative structure. Assortative structure is found in food webs when, e.g., node attributes related to environmental conditions correlate with interaction probability (e.g., fish swimming depth [37]). Disassortative structure is found in food webs when, for example, nodes differ in traits between trophic levels. In the RGG, the probability that a pair of nodes is connected is given by an attachment function parameterized by the Euclidean distance *d*(*i, j*) between a pair of nodes’ trait vectors, e.g., a decreasing function of distance in the case of assortative networks (see Materials & Methods).

Using synthetic networks to measure the performance of link prediction algorithms allows us to calculate the theoretical maximum prediction performance [39] in terms of the standard Area Under the ROC (Receiver Operating Characteristics) Curve (ROC-AUC) statistic [67], using the underlying probability of missing edges in the synthetic network model (see Note S1), and to measure performance systematically. We performed tests on these synthetic networks by dividing the observed links uniformly at random into 5 equal-sized groups and performing 5-fold cross validation for each algorithm’s link prediction performance. That is, in each iteration, we remove (hold out) a distinct 20% of the observed links from each food web (validation set), and we predict these missing links using models trained on the other 80% of links (training set, see Materials & Methods). Under this scheme, we systematiclaly evaluated three different versions of the stacked model for varying *ρ* (Fig. 3): a model with structural predictors only (“structure-only model”) (Table S1), a model with node attribute predictors only (“attribute-only model”) (Table S2), and a model with both types of predictors (“full model”). The structure-only model also included a subset of nearest neighbor predictors based entirely on network structure and the full model was the only model containing nearest neighbor predictors that combined node attributes with a node’s local topology (Table S3).

**FIG. 3.**
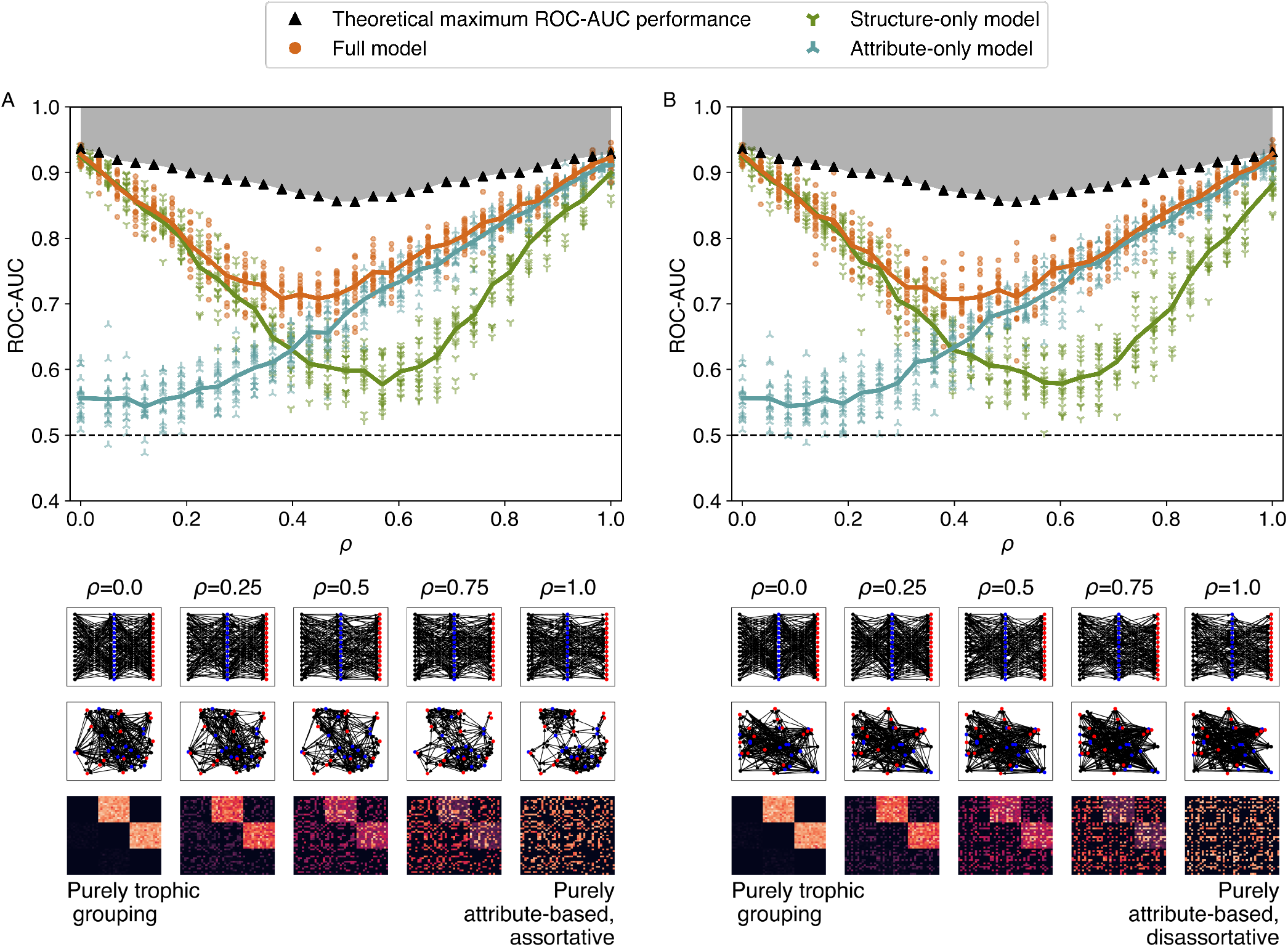
Link prediction on synthetic networks with known structure, using three stacked models: a structure-only model (47 predictors), an attribute-only model (20 predictors), and a ‘full’ model that includes both structure and attributes (71 predictors). Synthetic networks are a variable mix (see text) of purely trophic grouping structure without attributes (*ρ* = 0), and purely (A) assortative or (B) disassortative attribute-based connections without trophic groupings (*ρ* = 1). Thumbnails show examaple visualizations of mixture networks for specific choices of *ρ*. Main panels show the mean Area Under the ROC (Receiver Operating Characteristics) Curve (ROC-AUC) performance as a function of the mixing parameter *ρ* for 20 iid synthetic networks evaluated under 5-fold cross validation for each type of attribute pattern, along with the baseline ROC-AUC at 0.5 (dashed line) and the theoretical maximum performance at that *ρ* (black triangles). For both types of attribute pattern, the full model exhibits the best ROC-AUC performance at all values of *ρ*, matching the accuracy of the structure-only model when *ρ* = 0 and of the attribute-only model when *ρ* = 1. Moreover, at intermediate values of *ρ*, the full model performs better than either alternative model.

In this experiment, the performance of the structure model was highest when the synthetic network’s topology was drawn from the SBM anchor network (*ρ* ≈ 0.0), and nearly as high in the limiting case of drawing edges only from the RGG anchor network (*ρ* ≈ 1.0), in both assortative and disassortative cases. This latter behavior reflects the fact that spatial regularities in the generation of edges tends to induce specific topological patterns in network connectivity that the structure model can exploit. However, these patterns are distinct from those induced by group-based generation of edges, and hence the model’s performance was lowest when edges were drawn with nearly equal probability from the SBM anchor and the RGG anchor networks (*ρ* ≈ 0.6). In the limiting case of all edges drawn from the SBM anchor network, the structure model matched the theoretical maximum accuracy, and was only slightly suboptimal in the limiting case of all edges drawn from the RGG anchor network.

The performance of the attribute model was lowest when a majority of synthetic network’s topology was drawn from the SBM anchor network (*ρ <* 0.5), and was only slightly better than chance (ROC-AUC ≈ 0.56) when more than 80% of edges were drawn from the SBM anchor network. In contrast, the attribute model’s performance improved steadily as the fraction of edges drawn from the RGG anchor network increased above 20%. And, in the limiting case of all edges drawn from the RGG anchor network, the attribute model matched the theoretical maximum accuracy.

The full model performed well across all values of *ρ*, matching or exceeding the structure model for small values of *ρ* and matching or exceeding the attribute model for large values of *ρ*. The full model also exceeded the performance of both alternative models in the more difficult middle range of the mixing parameter *ρ* ≈ 0.5, demonstrating that the stacking model successfully learned how to best combine structure and attribute based predictors for missing link prediction, without knowing the underlying generative process.

### B. Performance on empirical food webs

We now evaluate the three stacked models—structure-only, attribute-only, and full models—using a large global database of empirical food webs [19, 20], which includes 290 networks across 5 ecosystem types (lakes, marine, streams, terrestrial aboveground, and terrestrial below-ground) with a common set of species traits as attributes for each node: log body mass, movement type, and metabolic type (Table I, see the Supplementary Information for details on data processing). We expected these traits to constrain interactions based on prior analyses of predator-prey interactions in this database [20].

**TABLE 1.** Species trait data included as node attributes in all 290 empirical food web studied here.

| Trait name | Type | Definition |
| --- | --- | --- |
| Metabolic type | Categorical (one-hot encoded) | 9 possible values: invertebrate, primary producer, ectotherm vertebrate, endotherm vertebrate, detritus, heterotrophic fungi, heterotrophic bacteria, dead organic material, other |
| Movement type | Categorical (one-hot encoded) | 7 possible values: walking, swimming, sessile, flying, other, floating, other_nonliving |
| Log<sub>10</sub> mass | Numeric (float) | Log base 10 of mean mass in grams of the population involved in the interaction |

For each food web, we divide the observed links uniformly at random into 5 equal-sized groups and perform 5-fold cross validation to assess each algorithm’s performance in realistic settings. We repeat this procedure for 5 independent iterations, thus averaging results over 25 iterations per food web. As with the synthetic networks, we measure algorithm performance using the ROC-AUC statistic, which provides a threshold- and scale-invariant measure of an algorithm’s ability to distinguish missing links (true positives) from non-edges (true negatives). In addition, we measure the Area Under the Precision-Recall Curve (PR-AUC) statistic. The PR-AUC provides a complementary measure to the ROC-AUC by emphasizing an algorithm’s ability to recover missing links (true positives) rather than simply its ability to correctly assign positives and negatives [84]. Across our evaluations, we examine performance relative to random baselines for both ROC-AUC and PR-AUC metrics. The baseline for the ROC-AUC is 0.5, while the baseline for PR-AUC differs by food web and is equal to the proportion of true positives (missing links) in the test set.

All three models produce mean ROC-AUC and PR-AUC scores far above the baselines for each empirical food web. Reflecting the versatility we observed on synthetic networks, the full model gives the highest mean performance across the 290 networks on both ROC-AUC and PR-AUC metrics (Fig. 4A,C), with mean ROC-AUC = 0.95 ± 0.06 and PR-AUC = 0.68 ± 0.2 (mean ± stddev), versus 0.94 ± 0.06 and 0.62 ± 0.2, respectively, for the structure-only model, and 0.88 ± 0.06 and 0.35 ± 0.2 for the attribute-only model. At the same time, however, on about 10% of the individual food webs for ROC-AUC and about 5% for PR-AUC, the attribute-only or structure-only models marginally outperformed the other models (Fig. 4B,D). Together, these results indicate that (i) both network structure and species traits are useful for predicting missing links in food webs, and (ii) the most accurate predictions are typically achieved by combining structural and trait information, i.e., they encode marginally different and complementary information about the existence of links, and (iii) model stacking is an effective way to learn from empirical observations alone the relative importance of network structure and species traits for making those predictions, without needing to know in advance how they correlate with links.

**FIG. 4.**
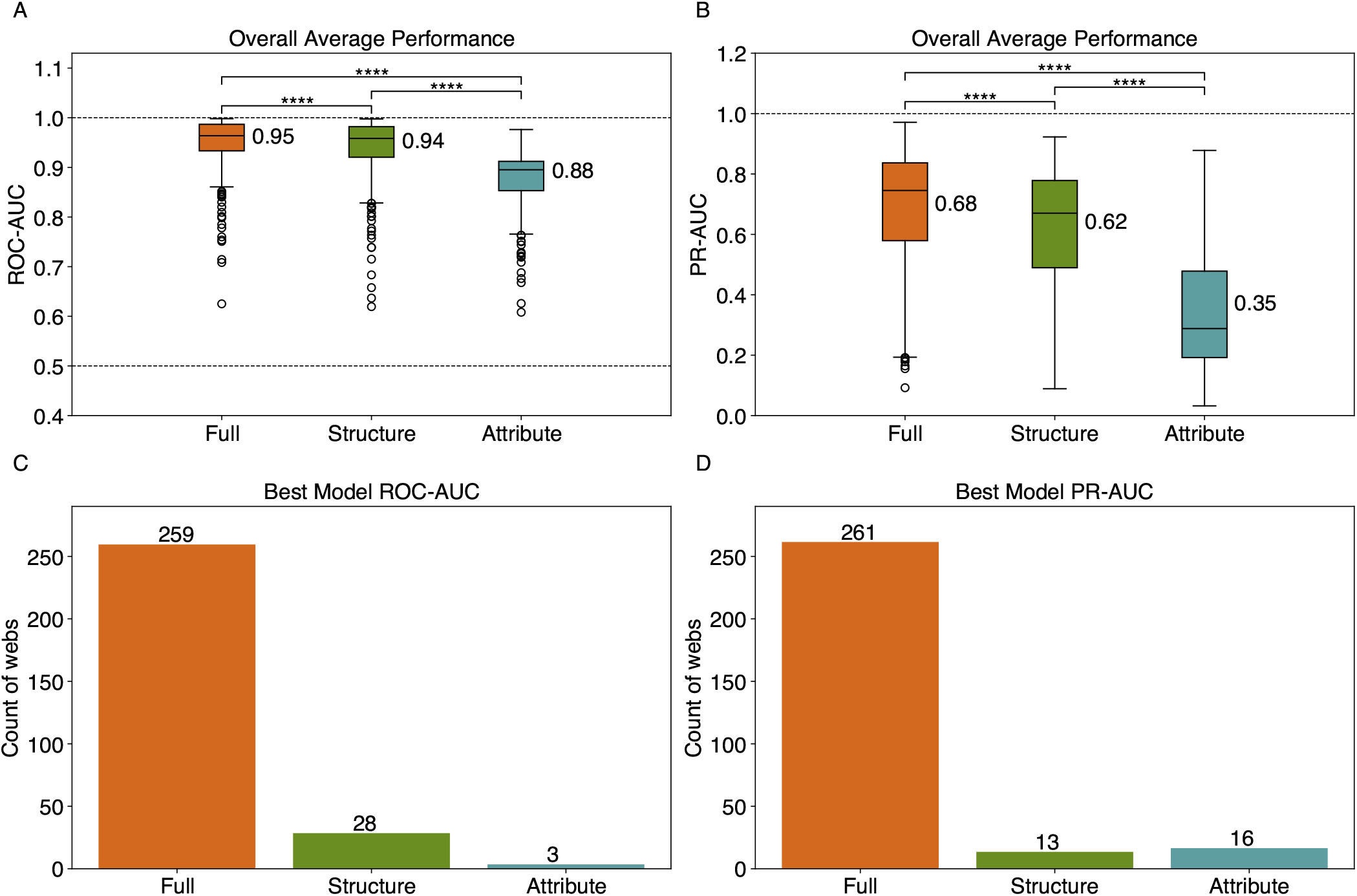
Link prediction performance on 290 food webs, for stacked models using structure-only predictors (‘structure’), attribute-only predictors (‘attribute’), and both (‘full’). Models are evaluated via (A) ROC-AUC and (B) PR-AUC metrics. Results are averaged for each food web across 5 independent iterations of evaluating across 5 unique folds (25 results per food web). Mean performance is displayed for each model across food webs. Significant differences in mean model performance based on false discovery rate (FDR) adjusted (Benjamini-Hochberg method [12]) within-subjects pair-wise two-sided t-tests are shown, where **** indicates a p-value*<*0.0001, along with (C,D) the respective counts of the number of food webs for which a particular model produced the best average score for each metric.

The food webs in our empirical corpus can be divided into five ecosystem types, which allows us to compare how link prediction performance varies with ecosystem characteristics. As with the aggregate results, we find that the full model is highly accurate at distinguishing missing links (true positives) from non-edges (true negatives). Moreover, the full model performs best on average in all five ecosystem types (Figs. 5 and S1), ranging from ROC-AUC = 0.99 *±* 0.01 on terrestrial below-ground food webs to 0.92 ± 0.06 on marine food webs. And, in each ecosystem, the structure-only model performed only marginally worse on average than the full model, with a larger step down in mean accuracy for the attribute-only model in each case. Marine food webs yield the smallest gap in ROC-AUC performance between the structure-only and attribute-only stacked models (ROC-AUC = 0.91 ± 0.07 vs. 0.88 ± 0.07, respectively), while terrestrial above-ground food webs yield the largest (ROC-AUC = 0.93 ± 0.08 vs. 0.79 ± 0.05, respectively). This modest variability in absolute prediction accuracy across the five ecosystem types suggests that ecosystem characteristics play an important but marginal role in determining the relative importance of structural and trait characteristics in whether a link exists or not.

**FIG. 5.**
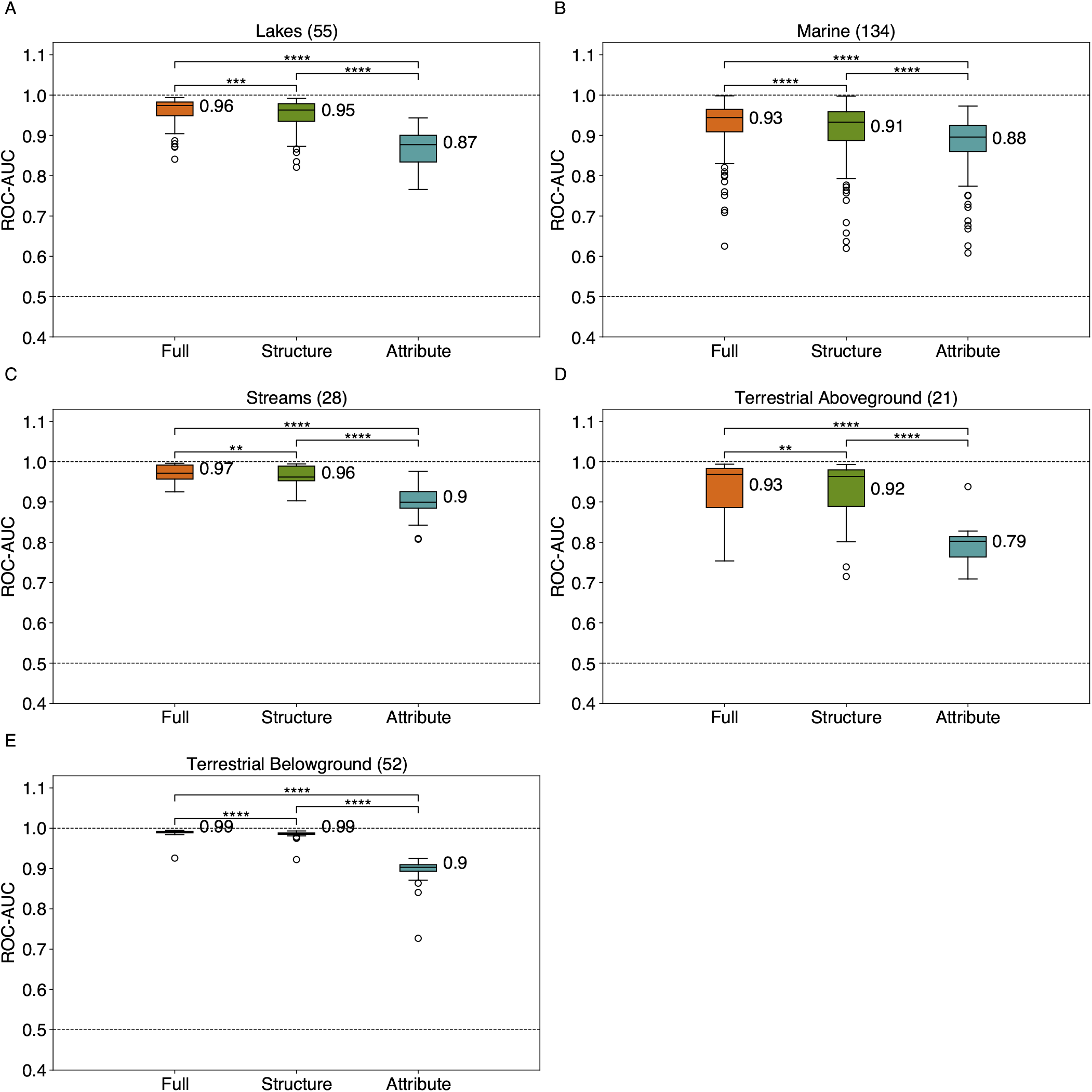
Link prediction performance by food web ecosystem type: (A) Lakes, (B) Marine, (C) Streams, (D) Terrestrial Aboveground, and (E) Terrestrial Belowground, for stacked models using structure-only predictors (‘structure’), attribute-only predictors (‘attribute’), and both (‘full’). The number of food webs in each ecosystem type is indicated in parentheses, mean ROC-AUC is displayed for each model across food webs, and significant differences in mean model performance based on false discovery rate adjusted (Benjamini-Hochberg method [12]) within-subjects pair-wise two-sided t-tests are shown, where **** indicates a p-value*<*0.0001, *** indicates a p-value*<*0.001, ** indicates a p-value*<*0.01, and * indicates a p-value*<*0.05.

An advantage of model stacking is that the supervised learning algorithm that sits atop the individual predictors can itself be inspected to learn which predictors the model identified as more or less useful in making accurate predictions. In our stacked models, Gini importance scores within the full models provide a quantitative measure of predictor utility (Fig. 6). In the full model, many of the structural predictors were among the most important, including two of the ecological predictors we adapted here (ecological preferential attachment, ecological Adamic/Adar index), with the ecological preferential attachment predictor achieving a higher relative importance. Predictors based on K-nearest neighbors (KNN), whether based entirely on structure or a combination of structure and node attributes, were also particularly helpful overall, as were predictors based on low rank approximations of the network, and some predictors encoding the centrality of node *j* (the consumer species). The log_10_ mass ratio between nodes was the most important attribute-based predictor. The top predictors varied across ecosystem types (Fig. S2), with a subset of predictors (ecological preferential attachment, KNN predictors, low rank approximation predictors) appearing as important across multiple ecosystem types, including when calculated using an alternative importance metric (permutation importance, Fig. S3).

**FIG. 6.**
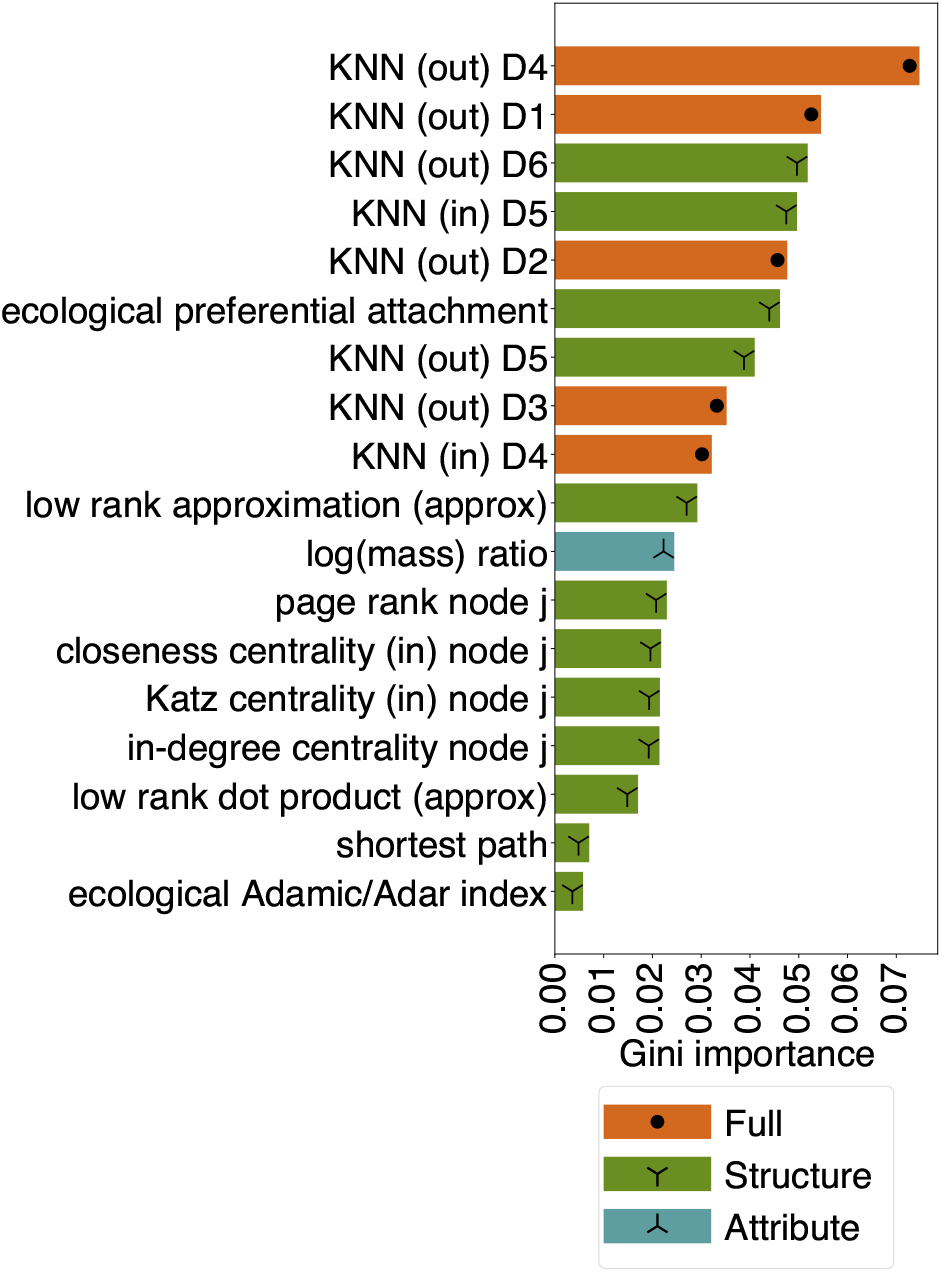
Top features by importance. Across 106 predictors in the full model, feature importance scores were averaged across folds and food webs. “Structure” predictors (out of 51) are based only on network structure, “attribute” predictors are based on node attributes (out of 47), and “full” predictors (out of 8) combine information about structure and node attributes in a single predictor. KNN predictors are based on K-Nearest Neighbors. The set of top predictors (18 total) was selected by taking the union of the top 10 predictors from each ecosystem type.

### C. Predictability of missing links depends on a food web’s characteristics

The large size of the empirical corpus of food webs allows us to investigate the determinants of a model’s predictive performance as a function of a food web’s characteristics, e.g., the fraction of species with missing body mass measures, whether parasites were removed or not, the distribution of species body masses, the food web’s size, and various summary statistics of the food web’s network structure. To do this, we first train a univariate regression that models missing link predictive performance (ROC-AUC or PR-AUC) as a function of food-web characteristics: (i) metadata and data processing (12 features, Table S4), (ii) global network topology (5 features, Table S5), and (iii) network assortativity (7 features, Table S6), adjusting for multiple comparisons. Empirical distributions of these feature are given in the Supplementary Information. We then inspect the regression coefficients to assess which characteristics of a food web correlate with more or less accurate predictions of missing links.

We find significant correlations between many network features and predictive performance for both ROC-AUC and PR-AUC (Fig. 7). For both ROC-AUC and PR-AUC, we found that all three models performed better on food webs that had lower proportions of nodes with an unclassified taxonomic level (Figs. S4A,S5A). This relationship was significant for all three models for the ROC-AUC. For the PR-AUC, this relationship was significant for the structure-only and full models, and for the attribute-only model was significant when controlling for the number of nodes in the network (Fig. S6B). We also found that the attribute-only model performed worse for networks with a higher proportion of nodes resolved at a taxonomic level higher than species for both metrics (Figs. S4B,S5B). Together, these results show that taxonomic resolution of the food webs was one of the factors that correlated with missing link predictive performance.

**FIG. 7.**
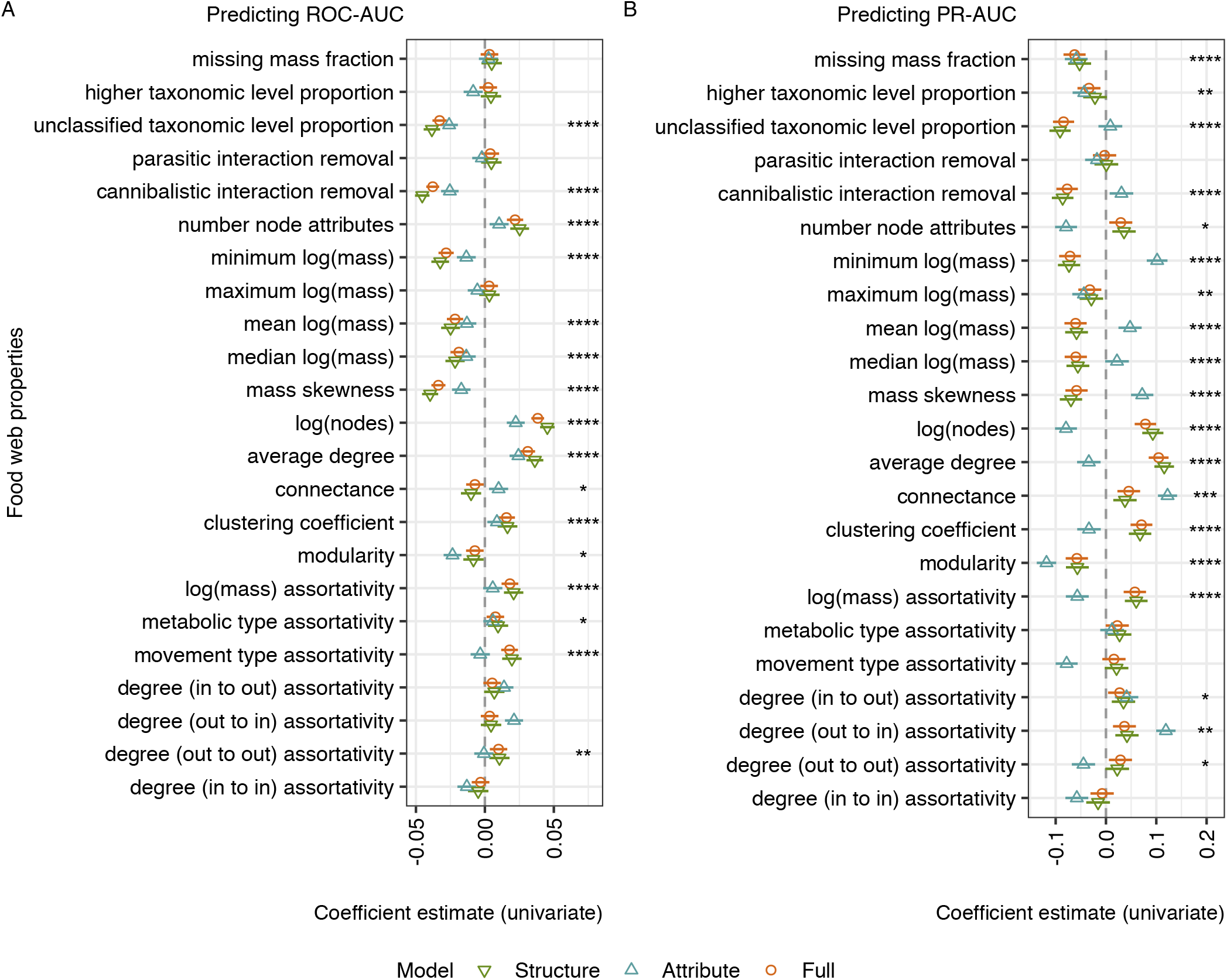
Correlates of missing link predictability. Regression coefficients in univariate linear regressions between food web properties and the mean (A) ROC-AUC and the (B) PR-AUC performance for missing link prediction. All food web properties were first *z*-score normalized so that the coefficients would be on comparable scales (un-scaled coefficient estimates are shown in Fig. S12). Whiskers show 95% confidence intervals and a vertical line at 0 represents neither a positive nor negative correlation. For the full model, **** indicates a *p <*0.0001, *** indicates a *p <*0.001, ** indicates a *p <*0.01, and * indicates a *p <*0.05 (FDR adjusted, Benjamini-Hochberg method [12]).

However, these significant trends related to taxonomic level were ecosystem type dependent (Figs. S7-S11). Notably, the observed trends with unclassified taxonomic level proportion were neutral or positive for all three models for both metrics looking only at stream food webs or only at terrestrial below-ground food webs. And, the observed overall trend with higher taxonomic level proportion did not hold when looking only at marine or terrestrial above ground food webs.

Looking at global topological metrics per network, we found that larger food webs (log number of nodes) generally had better link prediction performance, a trend that was significant for all three models for the ROC-AUC metric overall and for the structure-only and full models for the PR-AUC metric (Figs. S4C,S5C). However, there was a significant negative trend for the attribute-only model performance and log number of nodes for the PR-AUC metric, and the directions of the overall trends for the PR-AUC metric partially differed when looking at only stream, only lake, or only terrestrial belowground food webs (Figs. S7-S11). We found the same overall trends for average degree. However, these results differed when controlling for network size (Fig. S6), and in this case we observed that mean degree significantly correlated with better attribute-only model performance for both metrics, and only showed significant positive trends overall for the structure-only and full models for the PR-AUC metric. After controlling for network size, we found that connectance had a significant positive correlation with better performance for both metrics (Fig. S6). Together, these results indicate that link prediction was generally easier in cases where there was more link information available to the model both globally and locally.

We also found that link prediction performance was better for all models for food webs with less modularity (Figs. S4E,S5E), although this correlation did not hold looking only at lake or terrestrial aboveground food webs (Figs. S7-S11). For the 7 network assortativity features, we observed trends that were mixed across the features, ecosystem types, and models, particularly after controlling for network size (Fig. S6), indicating that assortativity did not display a universal trend with predictive performance.

## III. DISCUSSION

Food webs provide a broadly useful representation of the ecological complexity of species interactions. However, food webs are nearly always incompletely sampled because of the large number of potential interactions and the labor required to observe them, particularly rare interactions. Hence, more accurate methods for estimating missing links in a partially observed food web with commonly available species traits would improve the accuracy of data on species interactions, the efficiency of collecting it, and the utility of food web analyses and modeling. Here we evaluated the utility of model stacking—a state-of-the-art meta-learning technique for link prediction [39] that learns to combine multiple predictors into a single algorithm—for improving the accuracy of link prediction in food webs. Using this approach, we investigated the relative utility of species traits vs. species interactions for predicting missing links in food webs, how prediction accuracy varies with ecosystem type and network characteristics, and the relative utility of various individual link predictors.

Our broad analyses of synthetic food webs with known structure and of 290 real-world food webs indicates that species traits and observed network structure are both useful for predicting missing links—often because they are correlated—but, on average, structural predictors tend to produce more accurate predictions of missing links than do species traits. This result indicates that even when no trait data is available for nodes, structure-based methods can be used effectively to predict missing links in food webs. Of course, individual networks may be better explained by available traits alone or by structure alone, as has been found in previous work [114], and we find some evidence of this in our own results (Figs. 4, 5).

However, the most accurate predictions are generally obtained by combining traits and structural predictors within a single algorithm that can learn their relative importance in a particular food web, a result that is in alignment with prior work [31, 79, 89]. Moreover, across the empirical food webs studied here, we find that missing links tend to become more predictable when food webs are larger, have better taxonomic resolution, have higher connectance, and are less modular (Figs. 7,S6).

We also find that prediction accuracy and trait usefulness varies across ecosystem types: in the empirical food webs we studied, links in terrestrial belowground ecosystems are easiest to predict, while links in marine ecosystems are hardest to predict, and traits are least useful for prediction in terrestrial aboveground ecosystems. However, we had the fewest (21) food webs from terrestrial aboveground ecosystems, impeding generalizations. Although we do not test this possibility, the differences in methods used by different research teams to construct a food web may drive structural differences influencing predictor performance. For example, previous work has shown that there are distinct structural signatures in ecological networks based on construction methodology [18], which may influence which structural predictors are upweighted in the ensemble learned by model stacking.

Predicting missing links using species traits depends on what trait information is available for a given food web, and most empirical networks today include relatively few traits. For example, our analyses use just three traits (Table I). Substantial prior work has indicated that body mass plays a fundamental role in determining which edges exist in food webs as predators tend to be larger than prey and body size correlates with many species properties including locomotion, mortality, and abundance [15, 22, 31, 37, 40, 105, 113], and hence is likely to be broadly important in link prediction, regardless of ecosystem type [62, 100]. Our feature importance results also support this conclusion. At the same time, body mass is not a universal determinant of species interactions, a fact exemplified by parasitic and terrestrial herbivorous interactions, which tend to invert the usual direction of body mass’s influence, among other caveats [20, 21, 35, 51]. Feeding interaction prediction is typically improved by including other trait information along with body size [37, 89, 108, 110]. Hence, augmenting species interactions by collecting maximally detailed species trait information is likely to improve the prediction of missing links for both trait-based models [31, 83, 114] and the joint trait- and structure-based models we study here.

Previous work has also used information on species phylogenetic relationships as a proxy for missing traits [23, 24, 31, 41, 62, 72, 79, 85, 114], or inferred links based on species spatial co-occurrence [15, 103, 105, 107] (but see Refs. [7, 14, 29, 59] for discussions of methodological limitations). In particular, we do not consider any traits related to phenology or habitat overlap between species, which we would expect to constrain interactions. In adapting the model stacking approach to consider node attributes, we build upon particularly successful previous approaches that have used flexible machine learning methods to infer trait-matching rules [31, 62, 83, 114]. Machine learning methods will likely benefit from such additional trait information to learn useful trait-matching rules directly from data, shedding new light on the underlying ecology that structures species interactions.

Our results also reinforce and refine the relationship between species traits and network structure, showing variation across ecosystem types in the relative performance of the attribute-based model, as well as which attribute-based predictors are the most important. For example, our results show a smaller gap between the average performance of the attribute-based model and the structure-based model in marine food webs vs. terrestrial food webs. Collecting additional species traits beyond the three included in our modeling thus would be most important for missing link prediction in terrestrial food webs. We also see high feature importance for log_10_ mass ratio in marine food webs. Overall, these results may be in alignment with prior work showing marine food webs are more highly structured by body size than terrestrial food webs, a result with implications for theoretical models of food webs and stability analyses [65, 86, 95], and they highlight how link prediction with model stacking can be used to understand mechanistic differences in food web link formation between ecosystem types.

Stacking models like those considered here provide a further advantage by learning how to combine trait information with structural patterns to predict missing links, and we find that this combined learning approach produces superior performance compared to using structure alone or traits alone. In this way, we synthesize prior work on trait-matching in ecology with prior work on network structure [39, 106]. If network structural predictors improve missing link prediction beyond that possible using node attributes, this means that there are other latent rules driving link formation that are not captured by available node attribute data. Our results on both synthetically generated networks and empirical food webs show that a model stacking approach can be used to learn how to combine these predictors for a given network without knowing whether its link formation is driven by node attributes, by other latent rules, or by some combination of the two. Beyond improvements in prediction performance, model stacking can also facilitate analyzing the relative importance of different individual trait- or structure-based predictors [39, 43], in a way that is analogous to examining coefficients in a regression model. Insights about the features that drive better prediction performance can then be used to test specific ecological hypotheses or develop new ecological theories. For instance, a strong effect of structural predictors or phylogeny-based predictors, if included, relative to trait-based predictors, might suggest the existence of predator and prey adaptation not fully captured by the other species traits, such as exoskeleton hardness in coleopterans [108] or web-building vs. non web-building spider species [15].

Our feature importance results recapitulate ecological theory that the relative masses of two species and interactions between generalist consumers and generalist resources are important predictors of feeding links across all ecosystem types (Fig. 6). For instance, these patterns are key parts of the niche model for food web structure, in which species are ranked by a niche parameter, which is often associated with body size, and feed on species with lower values in this niche hierarchy within a range determined by a fundamental generality parameter [32, 40, 99, 111]. Our results also indicate the broad utility of KNN or “nearest-neighbor” predictors in food webs [10, 31, 114], which assume that closely related species will share traits and interaction partners, which is another phenomenon incorporated into models of food web structure to represent phylogenetic constraints on interactions [5, 26, 45, 52, 74, 97]. The existence of compartments of highly interacting nodes, for example due to habitat or seasonality constraints [5, 97], could also improve the performance of KNN predictors. Our results also show variation in the most important predictors in each ecosystem type, which supports investigating different generative models to best approximate food webs from different ecosystem types [65].

Our approach also has limitations. While we included many topological predictors in our stacked models, some of which have been used in prior work in ecological networks [30, 31, 88], there are other structure-based link prediction methods that we did not include as predictors within the ensemble. These include the probabilistic niche model [110], the allometric diet breadth model [82] (but see Ref. [4]), linear filtering [98], large scale models of grouping or hierarchical structure [5, 9, 28, 39, 44, 88], and matching-centrality latent trait models [88]. Future work could evaluate whether adding these as predictors within a stacking framework further improves predictions, or not, and whether using a different meta-learning model, such as a neural network approach [103] might improve performance.

While predictors based on low rank approximations were among the most important across food webs, we did not consider other link predictors based on learning a node embedding in a latent space—a technique that underlies many deep learning techniques [42, 56, 81, 102] including graph neural networks [46, 57, 109]— although recent work suggests that model stacking is often superior to these techniques for missing link prediction [39]. Instead, we focused on an ensemble of easy-to-compute topological and trait-based predictors that encode empirically-grounded aspects of food web structure, including hierarchical, clustering, and latent trait structure. Predictors like these have the advantage of not requiring complex or computationally expensive model fitting or node embedding procedures, and can be more easily interpreted in light of ecological mechanisms.

In the 290 food webs we analyzed here, we removed species interactions that were parasitic, cannibalistic, and repeated, following standard practice in the food webs literature. However, these interactions are increasingly believed to represent important information about ecosystems [60, 61], and a useful direction of future work may be to incorporate, and predict, the type of species interaction [71], perhaps using predictors for multi-level food webs (see Ref. [6]). For instance, some past work suggests that incorporating parasitic links can improve the prediction of missing non-parasitic interactions [53].

In addition, future work could explore more ecologically realistic, non-uniform patterns of link missingness, perhaps by modeling the characteristics that lead some links to be easier or harder to observe in the field, e.g., due to taxonomic or geographic biases of sampling [78], rather than simulating missing links by removing them uniformly at random [91, 104]. Recent work suggests that non-uniform missingness functions can lead to substantially different results compared to the uniform assumption [2, 104], even as model stacking tends to perform well even when links are missing in non-uniform ways [49].

In our predictive setting, we assumed that all nodes have the same set of attributes or species traits, which is not always the case, even for standard traits like body size. For example, in pre-processing the food web database, we had to make an assumption about how to assign placeholder body mass values for nodes like basal plants that lacked an observed value [114]. If more detailed trait data is collected, mismatches between sets of traits on some species vs. others will become more important to resolve. Finally, when evaluating missing link predictive performance we standardized on removing 20% of links. However, real-world ecological networks could be undersampled to far greater degrees, with the most extreme case being to construct a network from list of species and their traits alone.

In addition to understanding how state-of-the-art techniques like model stacking perform in such settings, two other use cases for link prediction algorithms represent promising future directions: (i) to guide the collection of data in the field, e.g., under an active learning framework [94], in which the model iteratively selects informative pairs of species to be observed by researchers, or (ii) to leverage more common data from some ecosystems to make predictions about interactions in another ecosystem, as under a transfer learning framework [24, 101, 102, 117] for which additional network-level features could be included [16].

Our results demonstrate the scientific utility of model stacking for predicting missing links in ecological networks, and their ability to learn how to combine both structural and trait-based information to improve predictions, regardless of whether traits exhibit an assortative or disassortative pattern with links. For future work in this area, an advantage of model stacking will be its easy extensibility: as new, theoretically grounded predictors are developed or as new data on traits or interactions becomes available, a stacked model can easily incorporate these new predictors in combination with existing techniques to produce more accurate predictions. Similarly, these techniques can adapt to more realistic models of missingness [49], can be used to predict future interactions in dynamic network settings [48], and may potentially be useful in guiding the collection of food web data in the field. We look forward to these many potential applications, and the benefits to ecological science they will bring.

## IV. METHODS AND MATERIALS

### A. Training and testing in model stacking for missing link prediction

In the missing link prediction problem, we assume that there is a true network *G* = (*V, E*) with a set of nodes *V* and a set of edges *E*; however, we only have access to an incomplete or observed network *G*^*′*^ = (*V, E*^*′*^), in which *E*^*′*^ ⊂ *E* is the incompletely observed set of edges. Let *X* = *V × V − E*^*′*^, the set of unconnected pairs of nodes in *G*^*′*^, denote the set of possible missing links, and define *Y* = *E − E*^*′*^ as the set of missing links (the positive class). We note that *Y* ⊂ *X*, and *X − Y* is the set of pairs of nodes that are unconnected in *G*, i.e., the true non-links (the negative class).

At a high level, the stacking model from Ref. [39] uses information from *G*^*′*^ to learn how to identify the pairs *i, j* ∈ *Y*, i.e., unconnected pairs in *G*^*′*^, that are in fact missing links *i, j* ∈ *X* relative to *G*. In order to train the stacking model, example missing links are produced from *G*^*′*^ via uniform removal to produce a training network *G*^*′′*^. Following Ref. [39], 1 ™ *α* is the probability that a link is removed to create this training dataset; thus, we remove (1 − *α*)*E*^*′*^ links. In our experiments, we set 1 − *α* = 0.2. For model training, the training dataset is composed of all unconnected pairs in *G*^*′′*^; removed links are the positive class and all non-links in *G*^*′*^ are taken as the negative class. Note that this negative class for training is noisy because it contains the true missing links from *G* (i.e., the set *Y*), but this effect on the model training is expected to be negligible for sparse networks [39]. In sparse networks like food webs, these classes are imbalanced with the negative class being much larger than the positive class. To better balance the classes before model training, we randomly up-sample with replacement the positive class, and for large networks we reduced the number of negative examples to 10,000 random examples of non-links to speed up model training [39].

A stacking model learns how to best combine the outputs of individual missing link prediction methods into an aggregate prediction for a given node pair. Features for the training data are generated for each candidate missing link by applying missing link prediction methods (detailed below). Each such method takes *G*^*′′*^ as input and produces a score or probability for each unconnected node pair *i, j*. In our stacking models, a supervised machine learning algorithm is used as the meta-learner and is trained on this dataset to differentiate between non-links (the negative class) and missing links (the positive class). This trained model is then used to make predictions for the test dataset (all pairs in *X*), with corresponding features generated from *G*^*′*^. The performance of the stacking model is then evaluated by comparing the ranking of pairs *i, j* ∈ *X* and whether they are missing links *i, j* ∈ *Y*. Here, we use a random forest [17] as the meta-learning model, with hyper-parameters chosen via 5-fold cross validation and optimized by selecting the best PR-AUC performance on average on the held-out fold, following advice in Ref. [84]. In our initial experiments, this choice slightly improved the downstream performance on food webs compared to using the F1 statistic, as in Ref. [39]. We note that there is some flexibility in the missing link prediction task regarding how one designs the validation, training, and test link sets [2], as well as in the choice of meta-level model and metric used for hyper-parameter optimization on the training set.

Importantly, food webs are typically represented as directed networks, with links pointing from resources (prey or primary producers) to consumers (predators, herbivores) [111], and sometimes have additional complexities, such as multiple interactions, self-loops, and edge weights. We consider food webs as simple directed networks, and remove all multiple interactions between species and all self-loops, which represent cannibalistic interactions. We leave a consideration of these complexities for future work. We note that for simple directed networks, the size of the set of potential m|issing links for a set of nodes *V* doubles from |*V*| (|*V* |*−*1)*/*2 as in Ref. [39] for undirected networks to |*V*| (|*V*| *−* 1), because *i, j* and *j, i* are independently potentially missing links. Hence, both directions are independently scored by the link prediction algorithm in our setting. This increases the size of the training set in *G*^*′′*^ from |*V* |(|*V* | − 1)*/*2 − |*E*^*′′*^| to |*V* |(|*V* |− 1)−|*E*^*′′*^|. Similarly, the size of the testing set in *G*^*′*^ increases from |*V* |(|*V*|−1)*/*2−|*E*^*′*^| to |*V* |(|*V* |−1)−|*E*^*′*^|.

### B. Synthetic network model parameters

We chose interaction probabilities for the SBM anchor networks and threshold values for the RGG anchor networks such that the median directed connectance value over 1000 generated networks matched that of a large database of empirical food webs: 0.125, 0.126, 0.127, and 0.127 for the food web database, SBM networks, assortative RGG networks, and disassortative RGG networks, respectively.

We generated SBM networks with 45 nodes and 3 equally-sized groups of 15 nodes with a high probability (*p* = 0.544) of interaction between nodes in groups 1 and 2, and between nodes in groups 2 and 3, and no probability of interaction (*p* = 0) among nodes between or within groups otherwise. This represents an extreme case of a food web with no omnivory, and is similar to model rectangular food webs with no omnivory used for modeling in prior work, e.g. as in Ref. [36]. (For more detailed discussion of functional groupings found in large food web datasets see Ref. [92]). The direction of the generated edges were chosen to replicate expected hierarchical structure in food webs by always pointing links from a lower group number to a higher group number; however, in a uniformly random 2% of cases (chosen to be similar to a large food web database, with a median value of 1.3%), edges were reciprocated (pointed in both directions).

To parameterize the RGG model, a random attribute vector was generated for each node consisting of four attributes: two numeric traits in the range [0,1] and two binary traits on {0, 1}. These traits were chosen to align with typical node attribute data for empirical food webs [19, 20] while also testing the ability to simultaneously consider both scalar and categorical traits in missing link prediction. The probability that two nodes are connected was then given by a simple step function, in which a pair of nodes is connected if the Euclidean distance between their attribute vectors was under a threshold for assortative networks [*d*(*i, j*) *<* 1.00375] or over a threshold for disassortative networks [*d*(*i, j*) *>* 1.425], and otherwise were not connected. Undirected edges were then converted to directed edges by randomly choosing an edge direction for each edge with equal probability, and again reciprocating edges in a uniformly random 2% of cases.

### C. Structural predictors

The structure-based models include 47 topological predictors for missing link prediction in food webs (see additional details in Table S1). Of these, 34 node and node-pair level topological predictors for undirected networks [39] were adapted for use in food webs. In addition to these, 18 were updated for directed networks by computing the same predictor but with a directed network rather than an undirected network as input:

- 6 predictors based on singular value decomposition, a strategy which has been shown to be good at predicting links in ecological networks [30, 98, 101, 102], using a directed version of the adjacency matrix: low rank approximation, low rank dot product, low rank mean, low rank approximation (approx), low rank dot product (approx), low rank mean (approx).
- 5 predictors based on shortest directed paths: shortest path, load centrality node *i*, load centrality node *j*, betweenness centrality node *i*, betweenness centrality node *j*.
- 4 predictors based on the number of local directed triangles: local clustering coefficient node *i*, local clustering coefficient node *j*, local triangles node *i*, local triangles node *j*.
- 3 page rank predictors, with a directed network as input: personalized page rank, page rank node *i*, page rank node *j*.
- 10 predictors were duplicated to calculate separate scores for both in- and out-directions:
- average neighbor in degree node *i*, average neighbor in degree node *j*, average neighbor out degree node *i*, average neighbor out degree node *j*;
  - closeness centrality (in) node *i*, closeness centrality (in) node *j*, closeness centrality (out) node *i*, closeness centrality (out) node *j*;
  - in-degree centrality node *i*, in-degree centrality node *j*, out-degree centrality node *i*, out-degree centrality node *j*;
  - eigenvector centrality (in) node *i*, eigenvector centrality (in) node *j*, eigenvector centrality (out) node *i*, eigenvector centrality (out) node *j* and
  - Katz centrality (in) node *i*, Katz centrality (in) node *j*, Katz centrality (out) node *i*, Katz centrality (out) node *j*.

Additionally, 6 topological predictors were adapted to our setting based on known topological properties of food webs. Food webs have directed links pointing from resource to consumer species and globally are hierarchically structured with links generally pointing in the direction of the flow of energy from lower to higher trophic levels. Food webs typically display many links between trophic levels and relatively fewer links within and across trophic levels, although links across a single trophic level can happen with omnivory. To account for this pattern of ecological organization, we adapted the preferential attachment (PA) link prediction method, which represents the intuition that two nodes with many links are more likely to share missing links, to an ‘ecological preferential attachment’ score (EPA) between nodes *i* and *j* by considering the product between the out-degree of species *i* and the in-degree of species *j*, thus predicting a higher likelihood of missing links between generalist resources and generalist consumers (see Eq. (1) and (2), where deg(*i*) represents the degree of *i*, deg^−^(*i*) represents the in-degree of *i* and deg^+^(*i*) represents the out-degree of *i*).

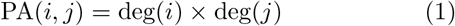

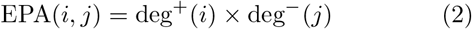

We also define the concept of a set of ecological common neighbors (ECNS, Eq. (4)) between a pair of species *i* and *j*. In undirected networks, the set of common neighbors (CNS, Eq. (3)) denotes the nodes that are connected to both *i* and *j*, and Γ(*i*) gives the neighbor set of node *i*. Common neighbor count (CN, Eq. (5)) predictors encode that we predict missing links by closing triangles of interactions [66]; however, this assumption is not appropriate for food webs because of their directed and hierarchical nature, and the rarity of loops [100]. Instead, the ‘ecological common neighbor’ count (ECN, Eq. (6)) predictor encodes that we expect to close omnivory motifs [27] (Fig. 1), where Γ^+^(*i*) represents the out-neighbor set of species *i* and Γ^−^(*i*) represents the in-neighbor set of species *i*.

We replace CNS with ECNS in 5 topological predictor calculations. For example, the Jaccard coefficient predictor (JC, Eq. (9)) which represents the number of common neighbors between two nodes (CN) divided by their number of total neighbors (TN, Eq. (7)) becomes the ‘ecological Jaccard coefficient’ predictor (EJC, Eq. (10)), in which we consider ecologically relevant common (ECN) and total (ETN) neighbors. The Leicht-Holme-Newman index [63], Adamic Adar index [1], and Resource Allocation index [116] are similarly updated and we add an additional predictor (ecological common neighbor scores) based on this concept (see Note S2). Finally, given the importance of trophic level for determining species interaction patterns and thus contextualizing other predictors, we added two indicators of approximate trophic level of a species calculated from the network structure (trophic level node *i* and trophic level node *j*) [64].

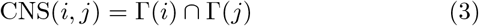

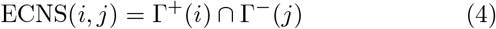

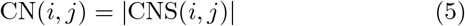

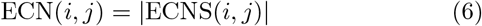

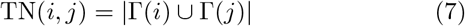

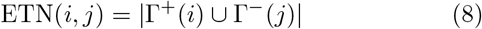

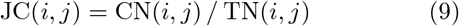

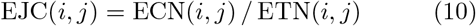

Some of these predictors, such as the Jaccard index, common neighbors, and degree product, have been previously explored for missing link prediction for food webs [88] and some have been noted as ecologically interpretable, for example the degree product can be interpreted as relating to the generality of the two species [79, 93].

### D. Attribute-based predictors

We additionally include predictors based on node attributes (species traits). We assume that all nodes have the same set of attributes *A*, which includes a subset of numeric attributes *N* and binary attributes *B, N ∩ B* = ø, *A* = *N* ∪ *B*. The synthetic and empirical food webs we consider vary in their attribute sets per node, and the number of predictors added for a given food web is a function of the size of these sets.

We add 2 *A* attribute features to each potential link simply by including attribute values for each of the nodes *i, j* in a pair, ordered based on the direction of the link between the two nodes (e.g., log(mass) of node *i*, log(mass) of node *j*), with categorical features first transformed into binary features via one-hot encoding.

Additionally, we add derived features based on the insight that many networks have assortative or disassortative structure based on node attributes (Fig. 2). In networks with assortative structure, interactions occur between nodes with similar attributes and in disassortative networks interactions occur between nodes with dissimilar attributes [75, 76]. These relationships can be incorporated into model stacking by adding distance metrics between the vectors of node attribute values. If |*N* | *>* 0, we include the Euclidean distance, Manhattan distance, cosine distance, and dot product between the numeric parts of the attribute vectors of node pairs (numeric attributes are first min-max normalized to the range [0, 1]). If |*B*| *>* 0, we also include the Hamming distance and Jaccard distance between the binary parts of the attribute vectors of node pairs. If | *A*| ≠ |*N* | and |*A*| = | | *B* (i.e., the nodes have a mix of numeric and binary attributes), then we also include the Euclidean distance, Manhattan distance, cosine distance, and dot product between the full attribute vectors.

Additionally, there may be some ratio between two numeric attributes that relates to the probability of a link. For example, trophic interactions have been shown to have typical log body mass ratios [20, 22]. We thus add the ratio between each of the numeric attributes of the two nodes, adding |*N*| additional predictors, taking into account the direction of a link (e.g., log(mass) ratio = log(mass)_*i*_ */* log(mass)_*j*_).

In total, we add 2 *A* attributes, up to 10 predictors measuring assortative or disassortative structure, and *N* attribute ratio predictors (Table S2). For example, with |*A* |= 9, |*N* | = 1 and |*B*| = 8, representing a typical case for the food webs we considered, we added 29 attribute-based predictors.

### E. Nearest-neighbor predictors

We additionally adapt the stacking model to include 12 K-Nearest Neighbor (KNN) predictors (Table S3). KNN predictors are based on the assumption that nodes that are similar have similar sets of interaction partners. KNN has performed well for link prediction in food webs in previous work [10, 31, 114]. KNN predictors are used to ‘recommend’ new interaction partners to a node. For example, in a food web with *K* = 2, a KNN predictor for species *i* would find the 2 most similar species to species *Ii* in the food web and then recommend prey species for species *i* from the prey sets of these two similar species with those prey found in both prey sets recommended first. The same approach can also be taken to recommend predators to prey.

We adapt KNN predictors for the stacking context by calculating for a node pair *i, j* the fraction of node *i*’s K-nearest neighbors in the food web that are in-neighbors (prey) of node *j*, as well as the fraction of node *j*’s K-nearest neighbors in the food web that are out-neighbors (predators) neighbors of node *i*. These predictors assume that if none of the nodes most similar to node *i* interact with node *j* and none of the nodes most similar to node *j* interact with node *i*, it is unlikely node *i* and node *j* interact whereas if all or most of the nodes similar to node *i* interact with node *j*, and the inverse, it is more likely they interact.

The nearness of neighbors can be determined by applying distance metrics to the attribute vectors or neighbor sets of two nodes. We used six distance metrics to produce 12 KNN predictors total based on both in- and out-directions (Fig. S13):

- D1: the Euclidean distance between the full normalized attribute vectors,
- D2: the Manhattan distance between the full normalized attribute vectors,
- D3: the Manhattan distance between the binary part of the attribute vectors,
- D4: the Jaccard distance between the binary part of the attribute vectors, following Ref. [31]
- D5: the Jaccard distance between in-neighbor sets (i.e., prey sets, also following Ref. [31])
- D6: the Jaccard distance between out-neighbor sets (i.e., predator sets).

D5 and D6 are calculated by subtracting the Jaccard similarity from 1 (Eqs. (11) and (12)). In cases of comparing two empty sets, for which this value is undefined, we set the Jaccard distance to 0 as we would expect two nodes that both don’t have prey (e.g., basal species) or two nodes that both don’t have predators (e.g., top predator species) to be similar. The predictors based on D5 and D6 only consider network structure, while the predictors based on D1, D2, D3, and D4 consider network structure and node attributes in combination. We implemented all KNN predictors with *K* = 3.

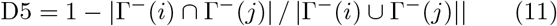

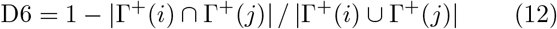

## V. ACKNOWLEDGEMENTS

The authors thank the Brose lab for help with the empirical data, A. Ghasemian for helpful conversations about stacking models for link prediction, and B. Singh, D.B. Larremore and E. Bradley for helpful discussions and feedback. This work was supported in part by the National Science Foundation Division of Ocean Sciences (NSF OCE 2049360) and the Eppley Foundation for Research. The stacking model code used here is adapted for use in directed, attributed, hierarchical networks from Ghasemian et al. (2020). The authors acknowledge the BioFrontiers Computing Core at the University of Colorado Boulder for providing High Performance Computing resources supported by BioFrontiers IT.

