## Supplementary material for "Predicting missing links in food webs using stacked models and species traits": Food_web_link_prediction_supplement_preprint_11_22_2024.pdf

### I. LIST OF SUPPLEMENTAL DATA FILES

- **Supplemental Data File 1.** Values filled in for nodes with movement type 'NA.'
- **Supplemental Data File 2.** Food web features across the database.
- **Supplemental Data File 3.** Summary statistics for food web features split by ecosystem type.

### II. NOTE S1. CALCULATING THEORETICAL ROC-AUC MAXIMUM PERFORMANCE

We contextualized our results on synthetic networks by calculating the theoretical maximum prediction performance in terms of the commonly used ROC-AUC statistic [31] for each value of $\rho$ using Monte Carlo sampling as in Ref. [15]. Mathematically, the ROC-AUC statistic is the probability that a higher score is assigned to a true positive (in this case, a missing link) than to a true negative (in this case, a non-link). At each value of $\rho$, we randomly sampled 10,000 pairs of true positives (edges) and true negatives (non-edges). For each pair, we then calculated the score as their probability of being connected from the known generative model (SBM, RGG, or a mixed SBM and RGG, all with an adjustment for the choice of edge direction). Our theoretical maximum ROC-AUC estimate was then the proportion of times out of these 10,000 samples that the true positive score was higher than the true negative score.

Further, we could examine partial theoretical maximum curves based on the SBM and RGG models individually by calculating the SBM and RGG probabilities of connecting edges separately. These partial curves are visualized for the case of an assortative RGG anchor network in Fig. S14.

### III. NOTE S2. CUSTOMIZING ECOLOGICAL TOPOLOGICAL PREDICTORS

We adapt the classic preferential attachment (PA, Eq. (1)) and common neighbor (CN, Eq. (5)) predictors to an ecological setting, to define the EPA (Eq. (2)), and ECN (Eq. (6)) predictors, where $\deg(i)$ represents the degree of node $i$, $\deg^{-}(i)$ represents the in-degree of node $i$, $\deg^{+}(i)$ represents the out-degree of node $i$, $\Gamma(i)$ represents the neighbor set of node $i$, $\Gamma^{+}(i)$ represents the out-neighbor set of node $i$, and $\Gamma^{-}(i)$ represents the in-neighbor set of node $i$. The EPA predictor assumes that interactions are more likely between generalist consumers and generalist resources. The ECN predictor is updated by substituting the common neighbor set (CNS) between nodes $i$ and $j$ (Eq. (3)) with the ecological common neighbor set (ECNS, Eq. (4), see Fig. 1).

$$\mathrm{PA}(i,j) = \deg(i) \times \deg(j) \tag{1}$$
$$\mathrm{EPA}(i,j) = \deg^{+}(i) \times \deg^{-}(j) \tag{2}$$
$$\mathrm{CNS}(i,j) = \Gamma(i) \cap \Gamma(j) \tag{3}$$
$$\mathrm{ECNS}(i,j) = \Gamma^{+}(i) \cap \Gamma^{-}(j) \tag{4}$$
$$\mathrm{CN}(i,j) = |\mathrm{CNS}(i,j)| \tag{5}$$
$$\mathrm{ECN}(i,j) = |\mathrm{ECNS}(i,j)| \tag{6}$$

We update the Leight-Holme-Newman index (LHN, Eq. (7)) [28] to the ecological Leight-Holme-Newman index (ELHN, Eq. (8)), by replacing its classic quantities with ecologically-oriented versions of those same quantities.

$$\mathrm{LHN}(i,j) = \mathrm{CN}(i,j)/\mathrm{PA}(i,j) \tag{7}$$
$$\mathrm{ELHN}(i,j) = \mathrm{ECN}(i,j)/\mathrm{EPA}(i,j) \tag{8}$$

We update the Adamic Adar index (AA, Eq. (9)) [1] to the ecological Adamic Adar index (EAA, Eq. (10)) in the same way, replacing its classic quantity CNS with the ecologically-oriented version ECNS. This predictor indicates that common neighbors of lower degree are potentially more impactful for predicting missing links.

$$\mathrm{AA}(i,j) = \sum_{w \in \mathrm{CNS}} \frac{1}{\ln(\deg(w))} \tag{9}$$

$$\mathrm{EAA}(i,j) = \sum_{w \in \mathrm{ECNS}} \frac{1}{\ln(\deg(w))} \tag{10}$$

We update the Resource Allocation index (RA, Eq. (11)) [49] to the ecological Resource Allocation index (ERA, Eq. (12)) in the same way.

$$\mathrm{RA}(i,j) = \sum_{w \in \mathrm{CNS}} \frac{1}{\deg(w)} \tag{11}$$

$$\mathrm{ERA}(i,j) = \sum_{w \in \mathrm{ECNS}} \frac{1}{\deg(w)} \tag{12}$$

We also add an additional score derived from all potential sets of directed common neighbors between two nodes ($\mathrm{ECN}_{\mathrm{score}}$, Eq. (16)).

$$\mathrm{C2}(i,j) = |\Gamma^{-}(i) \cap \Gamma^{+}(j)| \tag{13}$$

$$\mathrm{C3}(i,j) = |\Gamma^{+}(i) \cap \Gamma^{+}(j)| \tag{14}$$

$$\mathrm{C4}(i,j) = |\Gamma^{-}(i) \cap \Gamma^{-}(j)| \tag{15}$$

$$\mathrm{ECN}_{\mathrm{score}}(i,j) = \mathrm{ECN}(i,j) - [\mathrm{C2}(i,j) + \mathrm{C3}(i,j) + \mathrm{C4}(i,j)] \tag{16}$$

### IV. SUPPLEMENTAL FIGURES

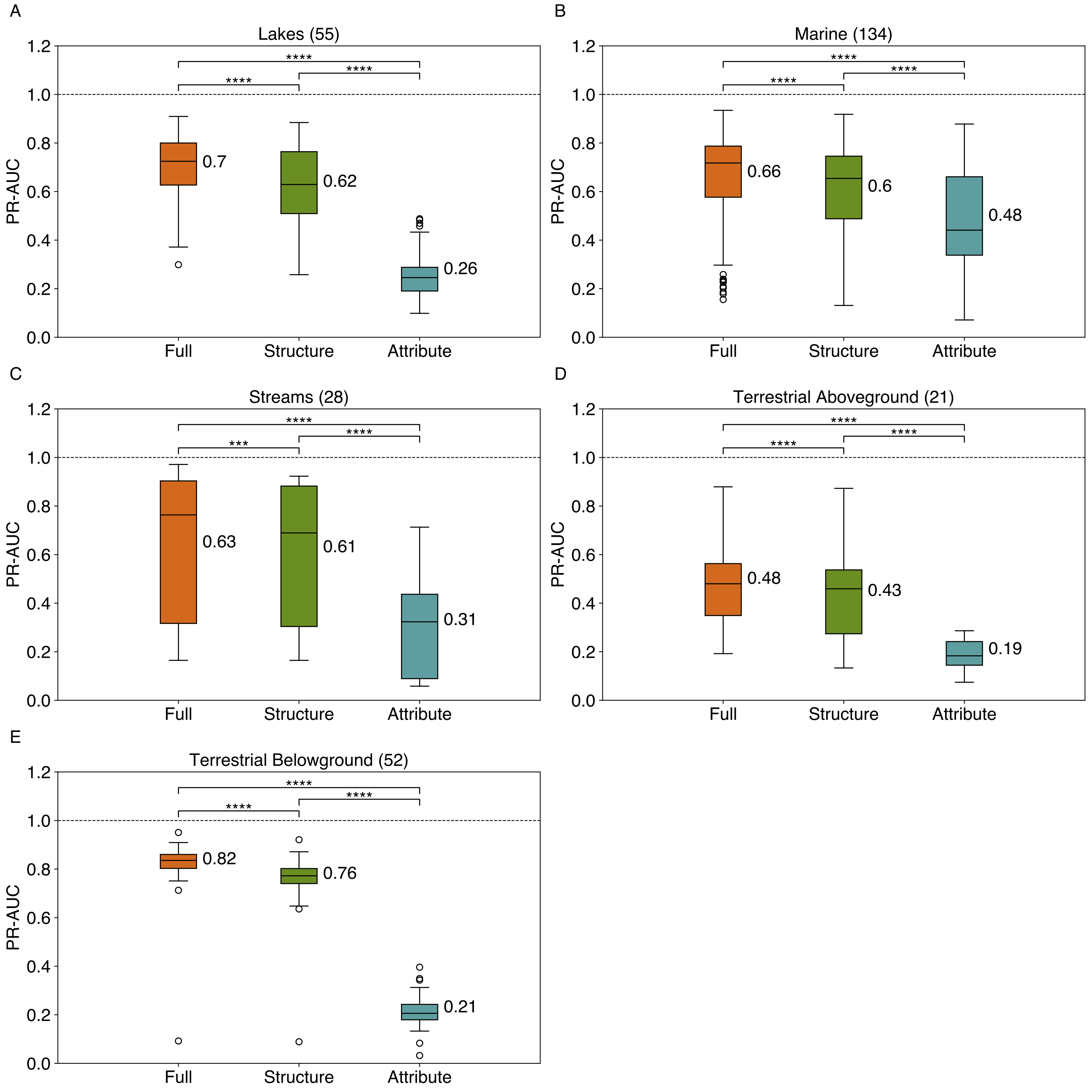


FIG. S1. Average link prediction performance for stacked models using structure-only predictors ('structure'), attribute-only predictors ('attribute'), and both ('full'), measured by Area Under the Precision-Recall Curve (PR-AUC), separated by ecosystem type: (A) Lakes, (B) Marine, (C) Streams, (D) Terrestrial Aboveground, and (E) Terrestrial Belowground. The number of food webs in each ecosystem type is indicated in parentheses, and significant differences in mean model performance based on false discovery rate (FDR) adjusted (Benjamini-Hochberg method [2]) within-subjects pair-wise two-sided t-tests are shown where **** indicates a p-value<0.0001 and ** indicates a p-value<0.01.

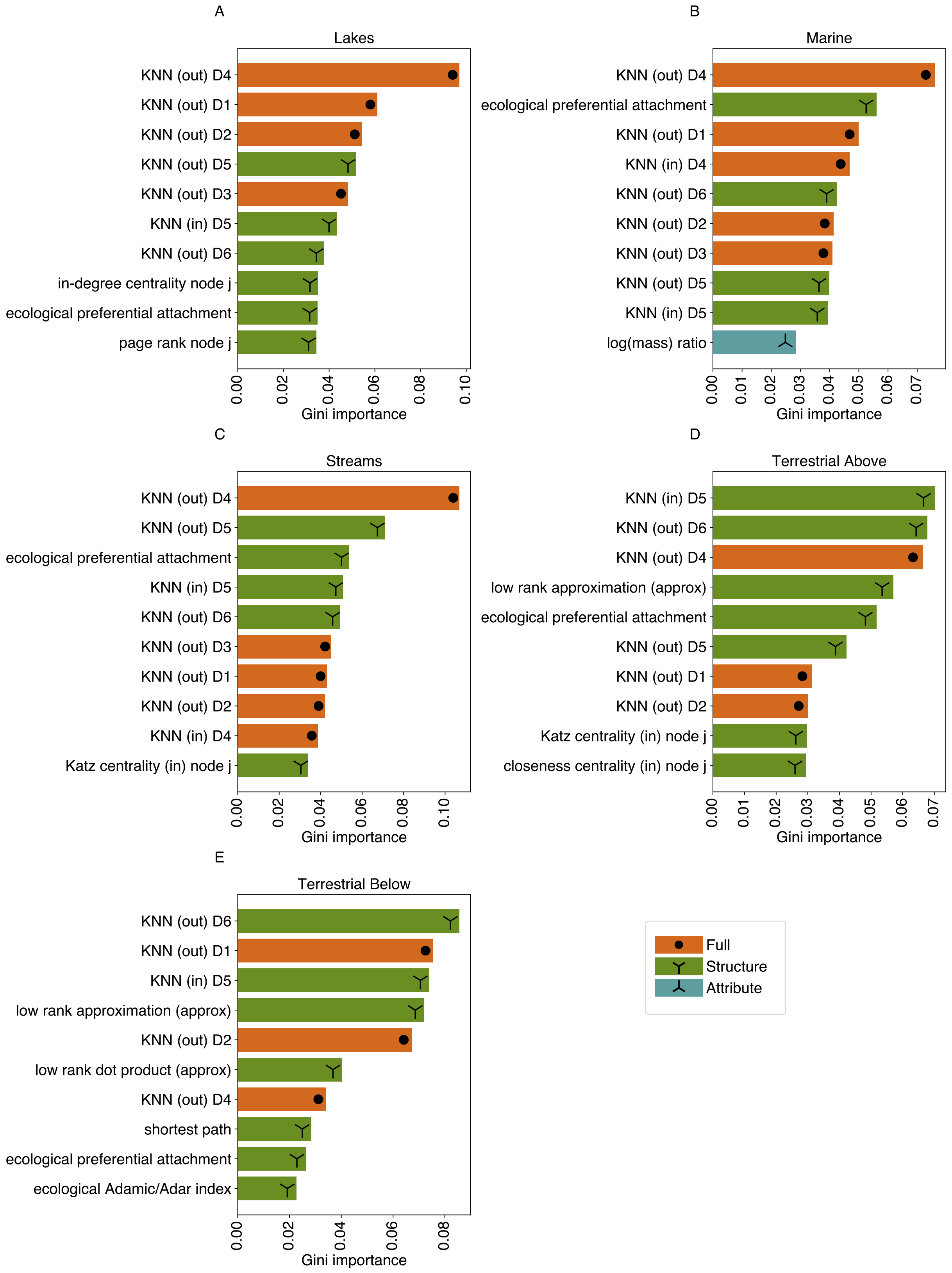


FIG. S2. Top 10 features ranked by Gini importance averaged across iterations and folds for food webs belonging to each ecosystem type: (A) Lakes, (B) Marine, (C) Streams, (D) Terrestrial Aboveground, and (E) Terrestrial Belowground. 'structure' predictors (out of 51) are based only on network structure, 'attribute' predictors are based on node attributes (out of 47), and 'full' predictors (out of 8) combine information about structure and node attributes in a single predictor.

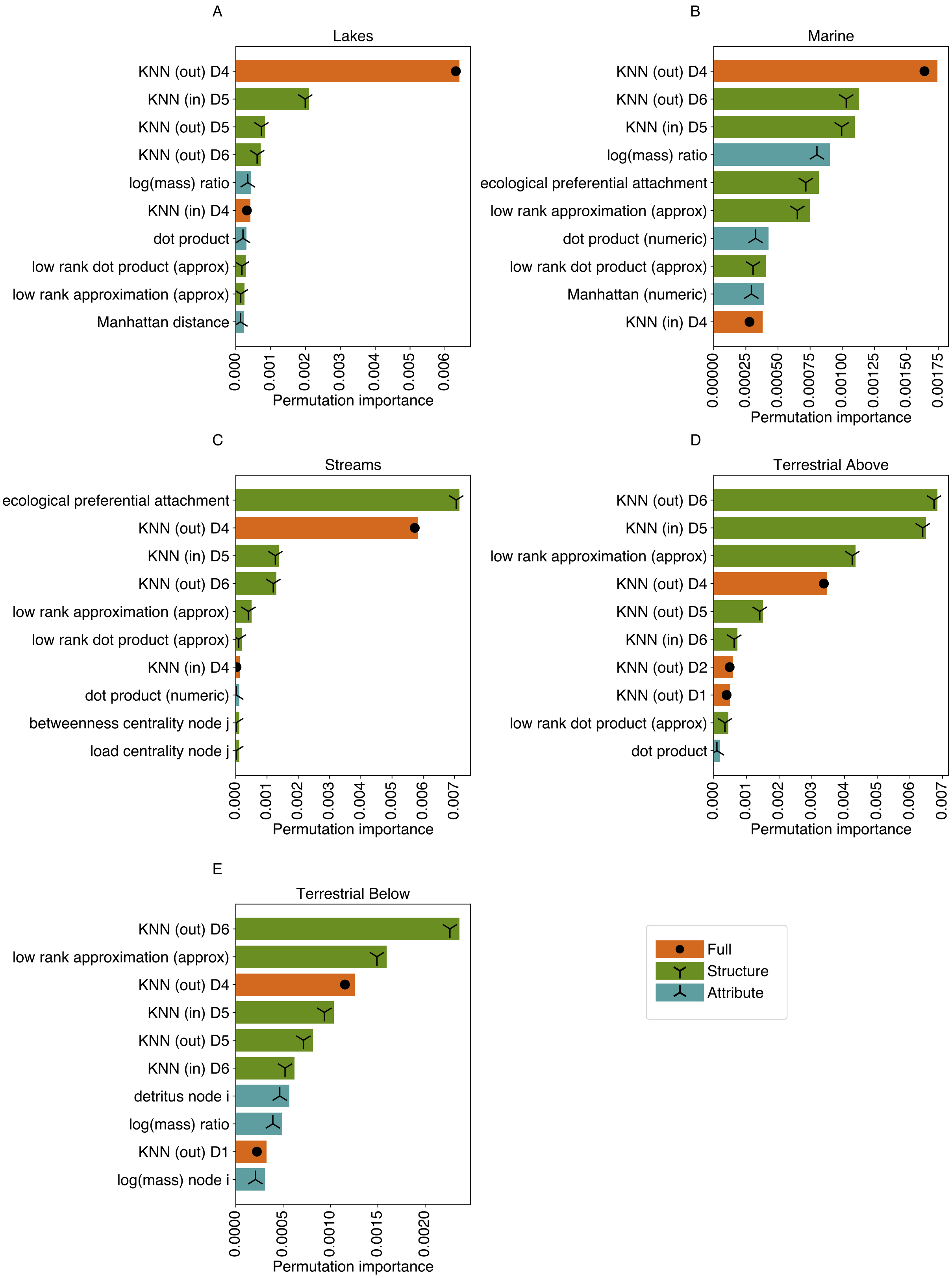


FIG. S3. Top 10 features ranked by permutation importance averaged across iterations and folds for food webs belonging to each ecosystem type: (A) Lakes, (B) Marine, (C) Streams, (D) Terrestrial Aboveground, and (E) Terrestrial Belowground. 'Structure' predictors (out of 51) are based only on network structure, 'attribute' predictors are based on node attributes (out of 47), and 'full' predictors (out of 8) combine information about structure and node attributes in a single predictor.

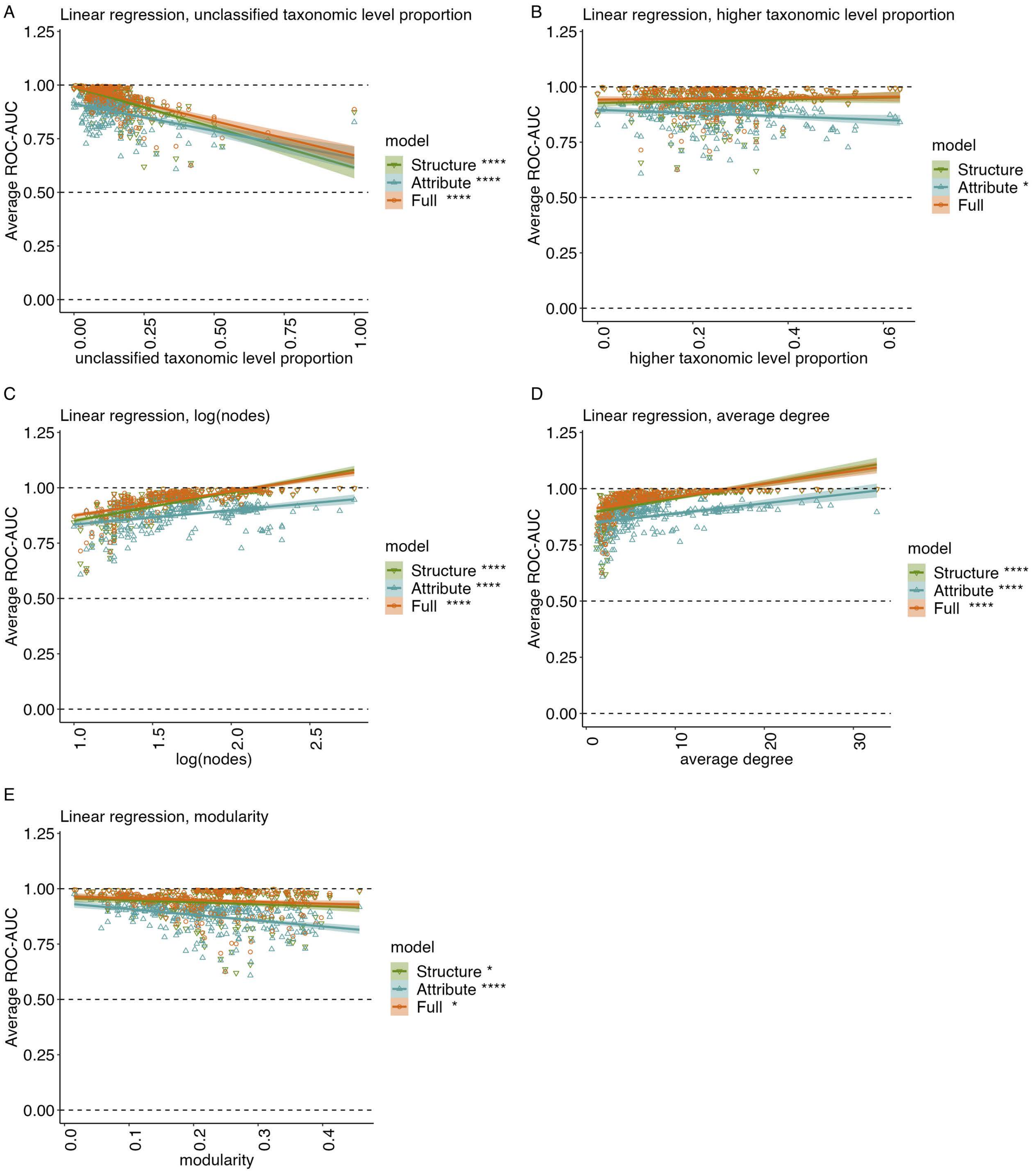


FIG. S4. Performance of the structure-only, attribute-only, and full models contextualized by select dataset characteristics. Average ROC-AUC results by (A) unclassified taxonomic level proportion, (B) higher taxonomic level proportion, (C) log(nodes), (D) average degree, and (E) modularity. Significance of coefficients in univariate linear regressions fit to these relationships are indicated by level ($p \leq 0.0001$: ****, $p \leq 0.001$: ***,$p \leq 0.01$: **, $p \leq 0.05$: *, FDR adjusted). Horizontal dashed lines indicate the threshold for better than random predictive performance (ROC-AUC=0.5) and the maximum possible performance (ROC-AUC=1).

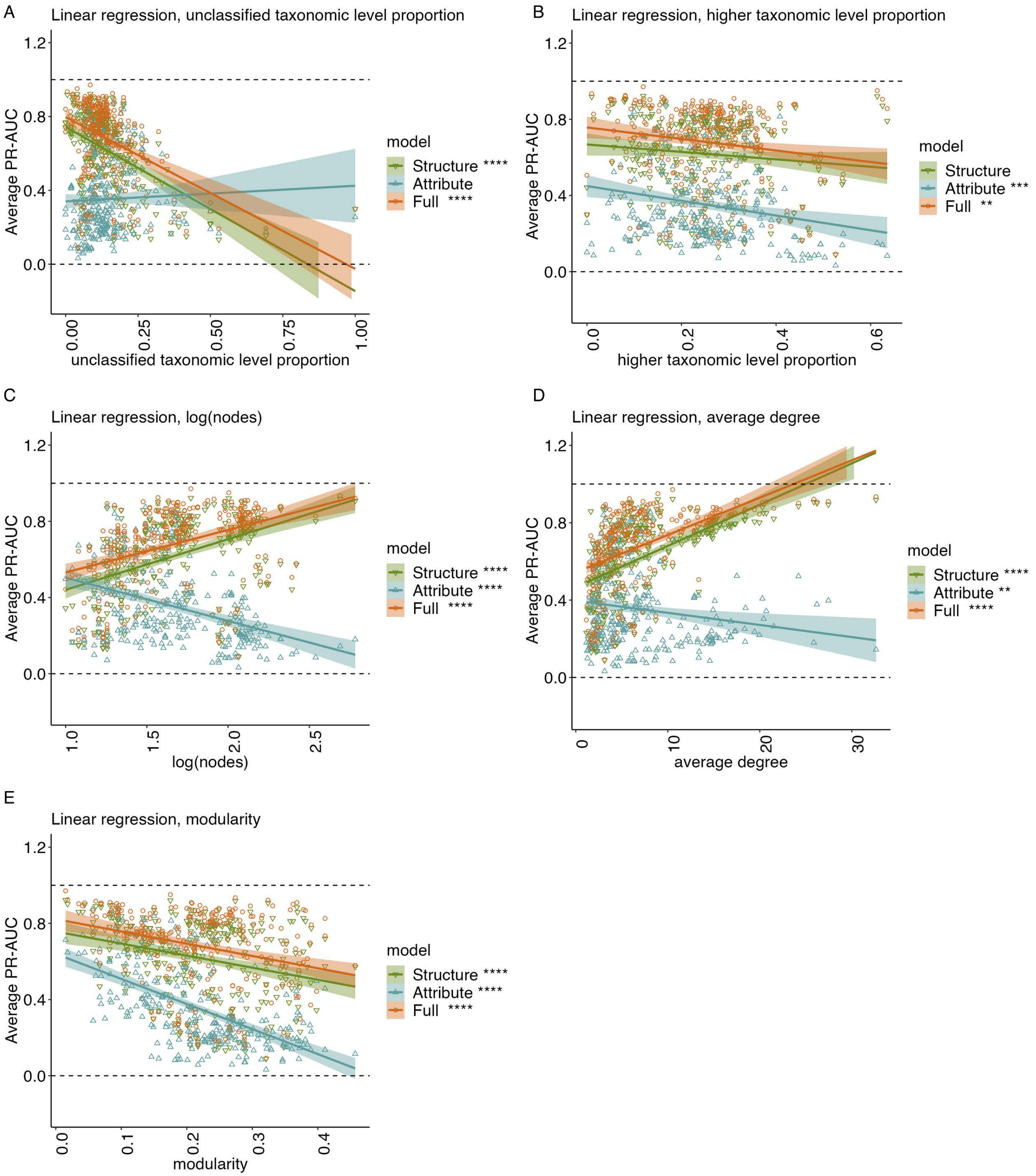


FIG. S5. Performance of the structure-only, attribute-only, and full models contextualized by select dataset characteristics. Average PR-AUC results by (A) unclassified taxonomic level proportion, (B) higher taxonomic level proportion, (C) log(nodes), (D) average degree, and (E) modularity. Significance of coefficients in univariate linear regressions fit to these relationships are indicated by level ($p \leq 0.0001$: ****, $p \leq 0.001$: ***,$p \leq 0.01$: **, $p \leq 0.05$: *, FDR adjusted). Horizontal dashed lines indicate the minimum possible performance (PR-AUC=0) and the maximum possible performance (PR-AUC=1).

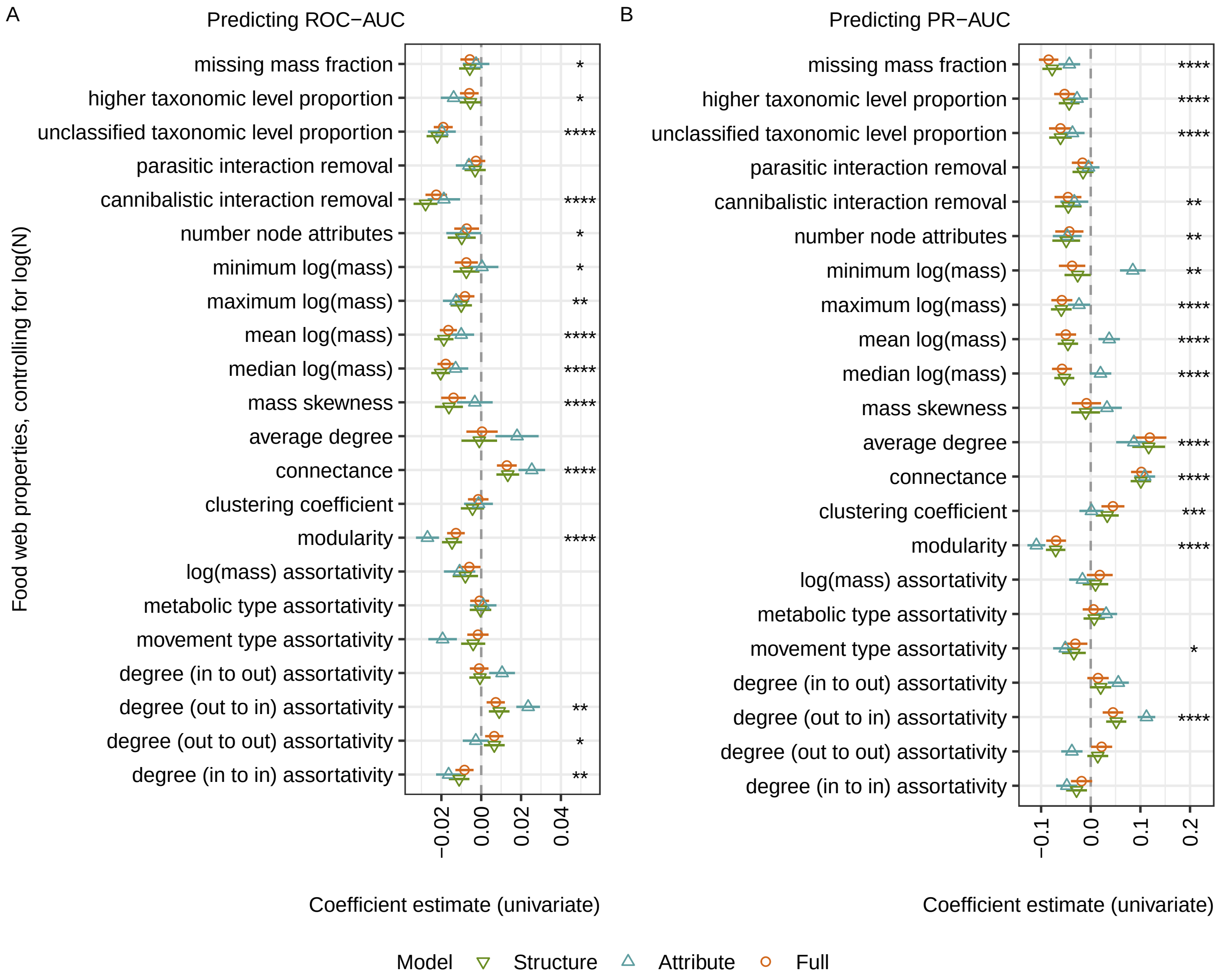


FIG. S6. Regression coefficients in univariate linear regressions between food web properties and (A) ROC-AUC and (B) PR-AUC performance for missing link prediction, when the log of the number of nodes in the food web is also included as a covariate. All food web properties were first z-score normalized. Whiskers show 95% confidence intervals and a vertical line at 0 represents neither a positive nor negative correlation. For the Full model, **** indicates a p-value<0.0001, *** indicates a p-value<0.001, ** indicates a p-value<0.01, and * indicates a p-value<0.05 (false discovery rate adjusted, Benjamini-Hochberg method [2]).

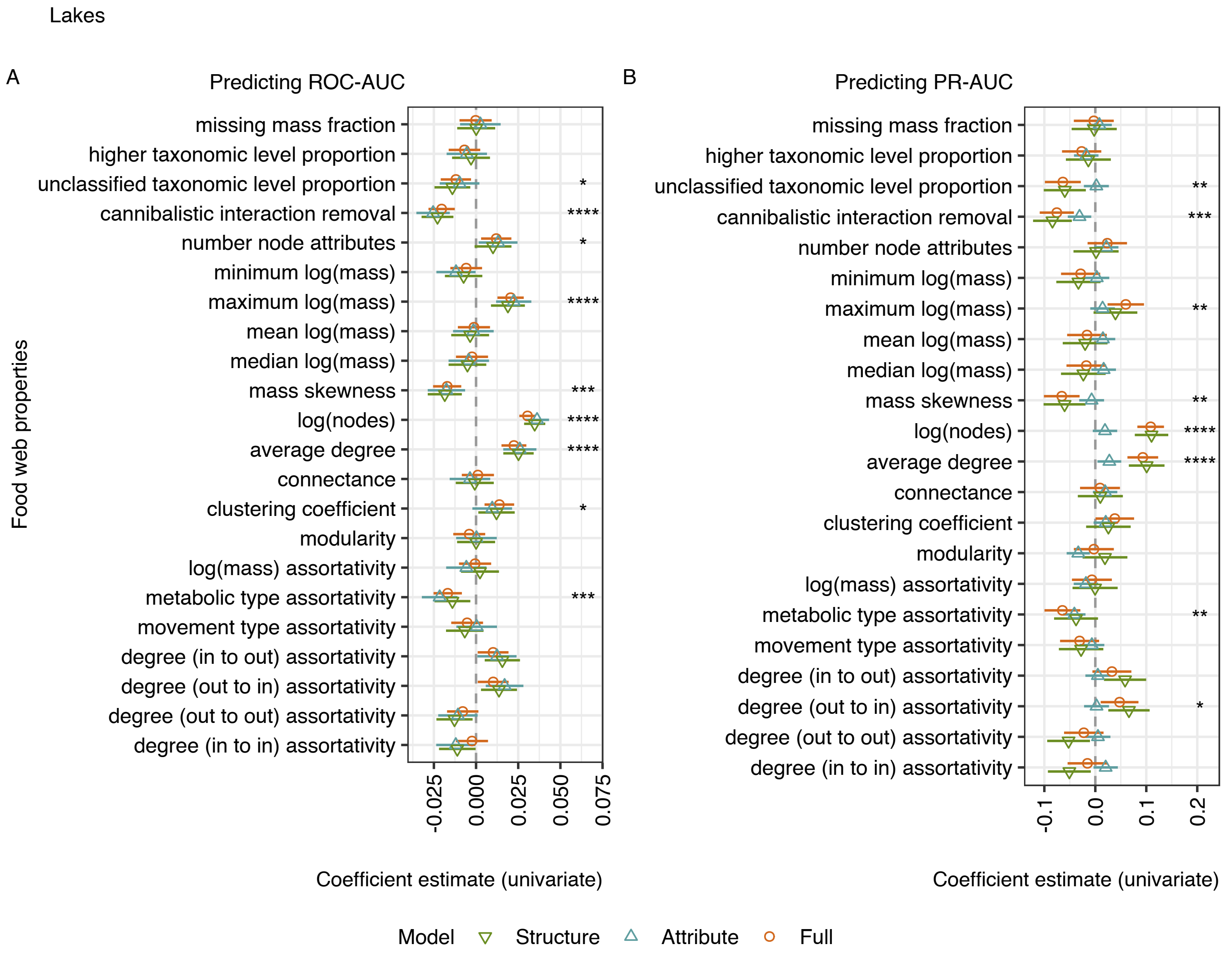


FIG. S7. Regression coefficients in univariate linear regressions between food web properties and (A) ROC-AUC and (B) PR-AUC performance for missing link prediction, for only lake food webs. For the Full model, **** indicates a p-value<0.0001, *** indicates a p-value<0.001, ** indicates a p-value<0.01, and * indicates a p-value<0.05 (false discovery rate adjusted, Benjamini-Hochberg method [2]). Parasitic interaction removal is not included as no lake food webs had parasitic interactions removed.

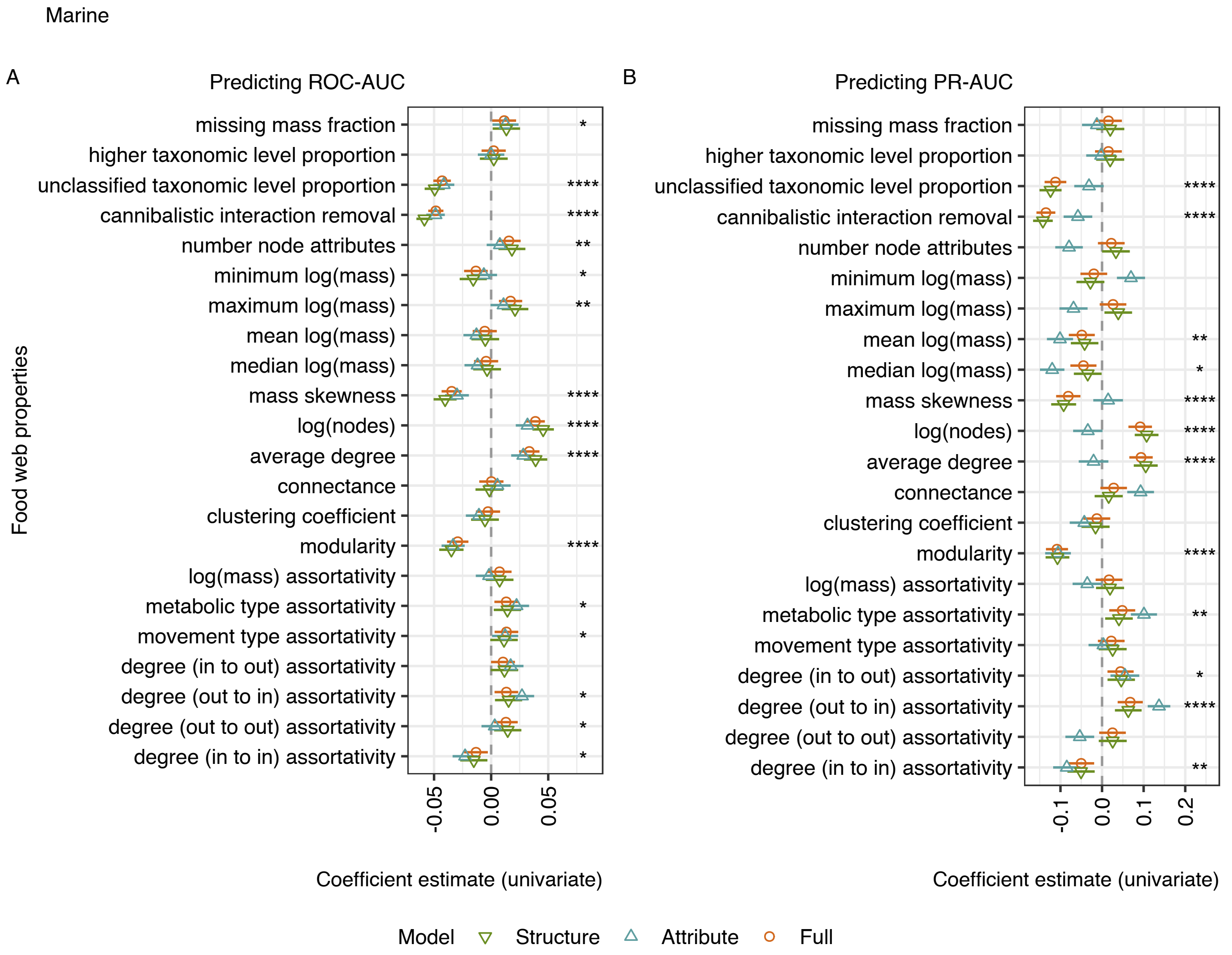


FIG. S8. Regression coefficients in univariate linear regressions between food web properties and (A) ROC-AUC and (B) PR-AUC performance for missing link prediction, for only marine food webs. For the Full model, **** indicates a p-value<0.0001, *** indicates a p-value<0.001, ** indicates a p-value<0.01, and * indicates a p-value<0.05 (false discovery rate adjusted, Benjamini-Hochberg method [2]). Parasitic interaction removal is not included as no marine food webs had parasitic interactions removed.

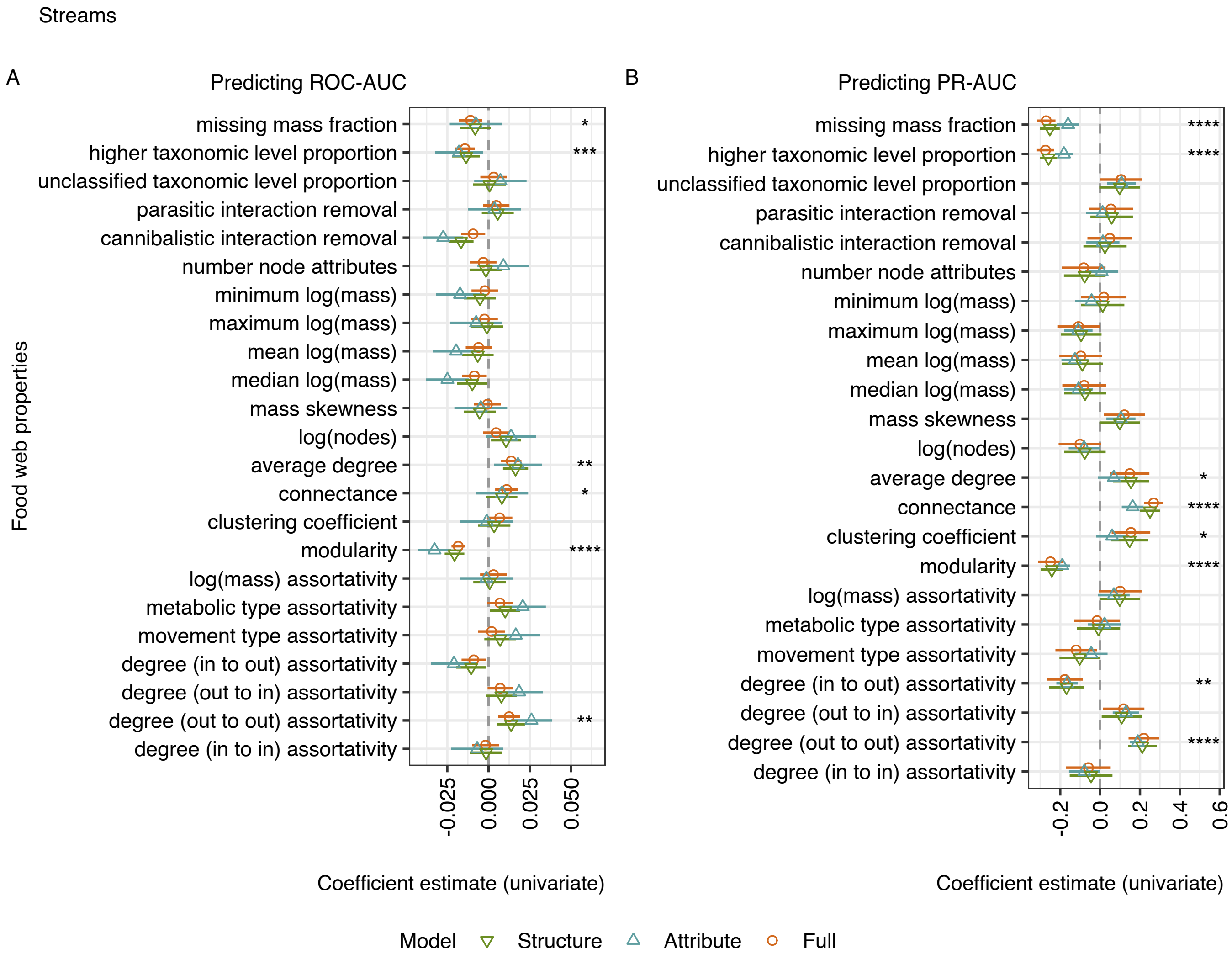


FIG. S9. Regression coefficients in univariate linear regressions between food web properties and (A) ROC-AUC and (B) PR-AUC performance for missing link prediction, for only stream food webs. For the Full model, **** indicates a p-value<0.0001, *** indicates a p-value<0.001, ** indicates a p-value<0.01, and * indicates a p-value<0.05 (false discovery rate adjusted, Benjamini-Hochberg method [2]).

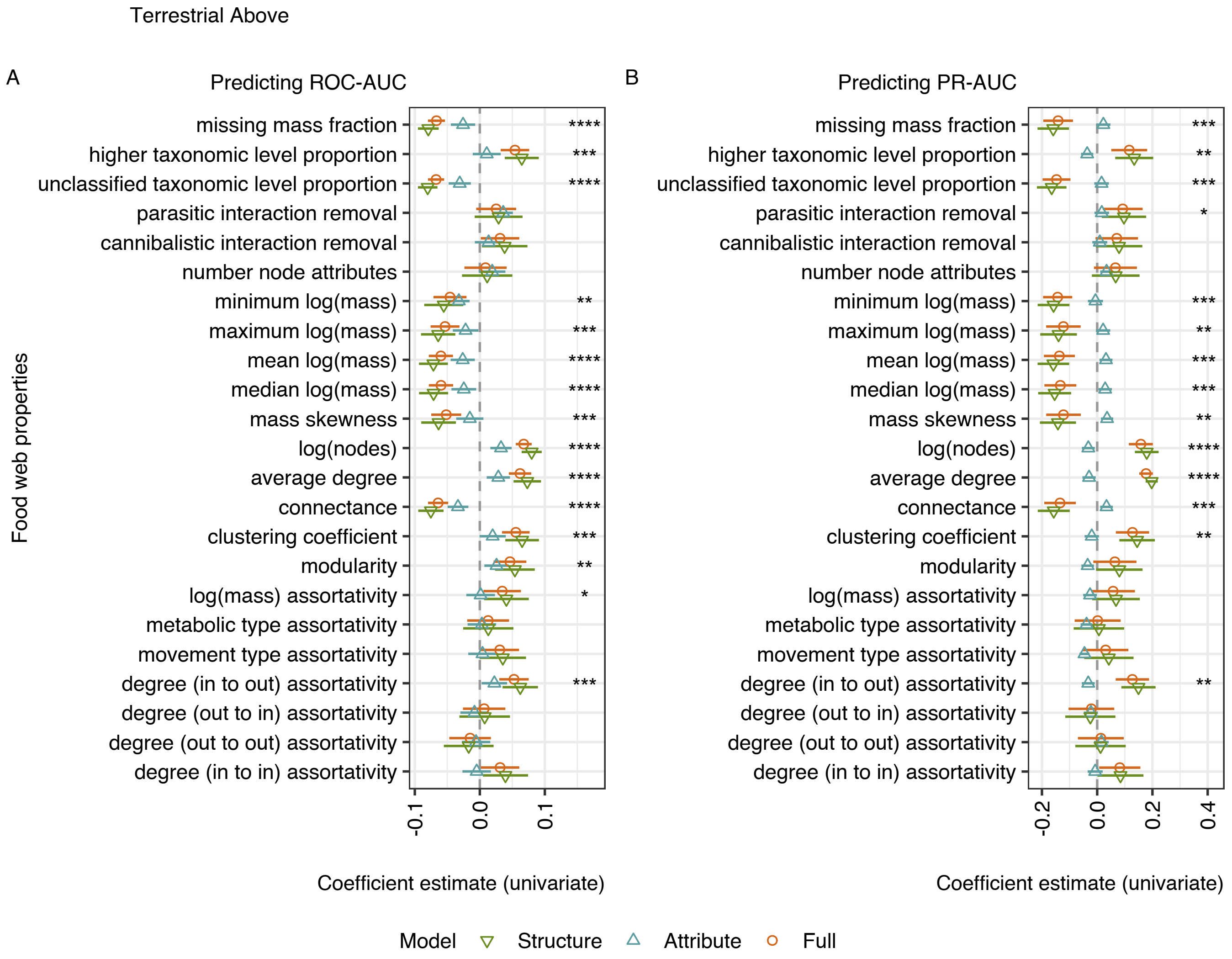


FIG. S10. Regression coefficients in univariate linear regressions between food web properties and (A) ROC-AUC and (B) PR-AUC performance for missing link prediction, for only terrestrial aboveground food webs. For the Full model, **** indicates a p-value<0.0001, *** indicates a p-value<0.001, ** indicates a p-value<0.01, and * indicates a p-value<0.05 (false discovery rate adjusted, Benjamini-Hochberg method [2]).

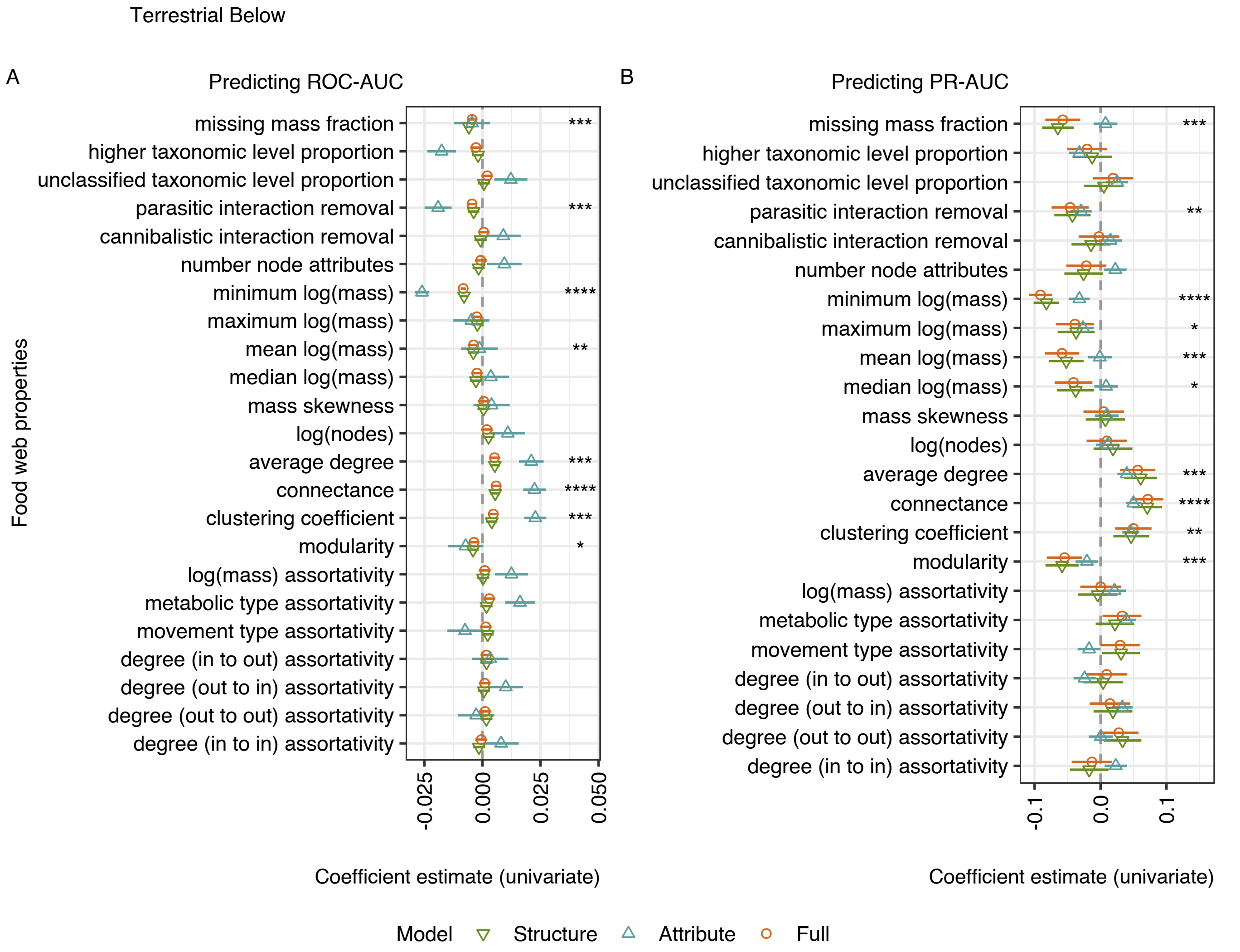


FIG. S11. Regression coefficients in univariate linear regressions between food web properties and (A) ROC-AUC and (B) PR-AUC performance for missing link prediction, for only terrestrial belowground food webs. For the Full model, **** indicates a p-value<0.0001, *** indicates a p-value<0.001, ** indicates a p-value<0.01, and * indicates a p-value<0.05 (false discovery rate adjusted, Benjamini-Hochberg method [2]).

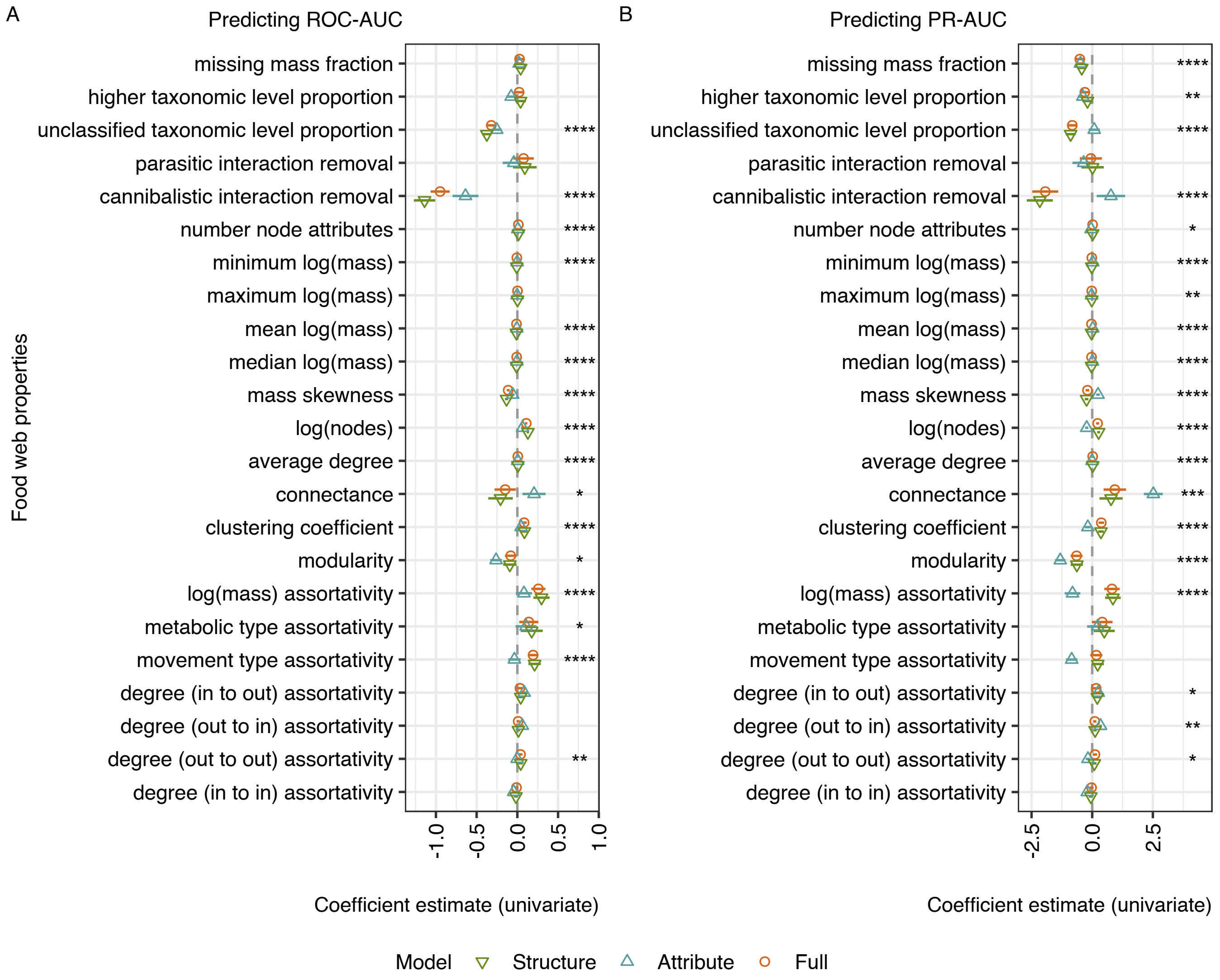


FIG. S12. Coefficient estimates in univariate linear regressions between food web properties and (A) ROC-AUC and (B) PR-AUC performance for missing link prediction, un-scaled. Whiskers show 95% confidence intervals and a vertical line at 0 represents neither a positive nor negative correlation. For the Full model, **** indicates a p-value<0.0001, *** indicates a p-value<0.001, ** indicates a p-value<0.01, and * indicates a p-value<0.05 (false discovery rate adjusted, Benjamini-Hochberg method [2]).

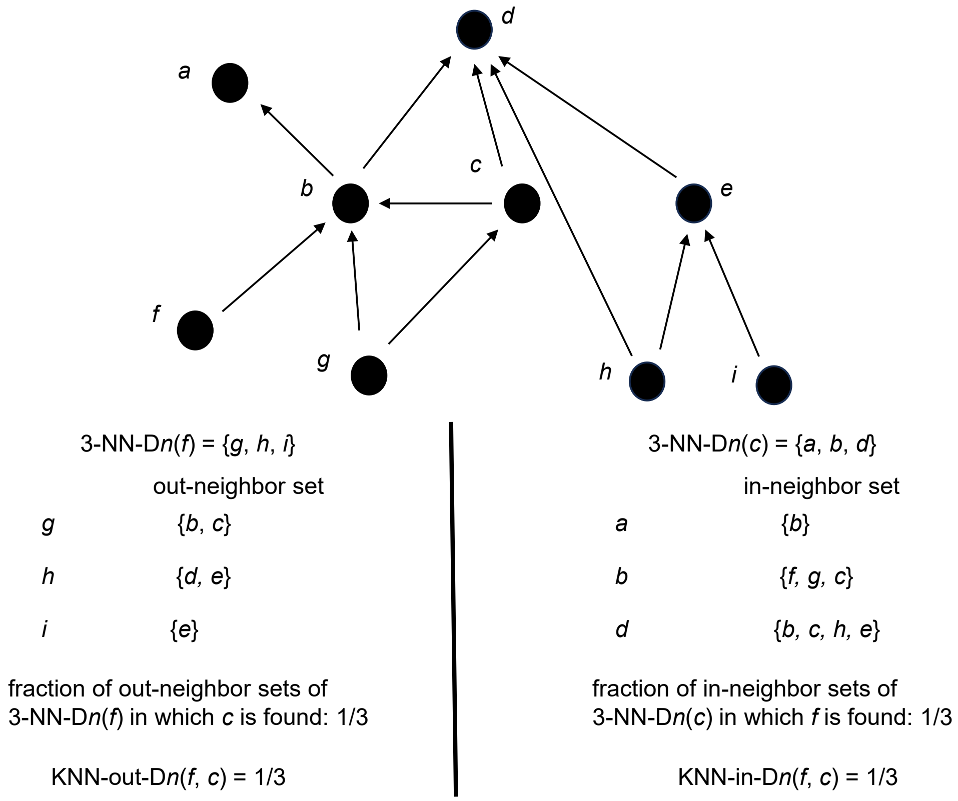


FIG. S13. KNN predictors. With $K = 3$, two KNN predictors are evaluated for in and out directions for each distance metric $D_n$. Here they are evaluated for the directed node pair $(f, c)$ in an example food web with a hypothetical distance metric $D_n$. After the 3 nearest neighbors are identified for each of $f$ and $c$ (3-NN-$D_n(f)$, 3-NN-$D_n(c)$), the out and in-neighbor sets are evaluated respectively to calculate a missing link prediction score based on how often $c$ or $f$ interact with the 3 nearest neighbors.

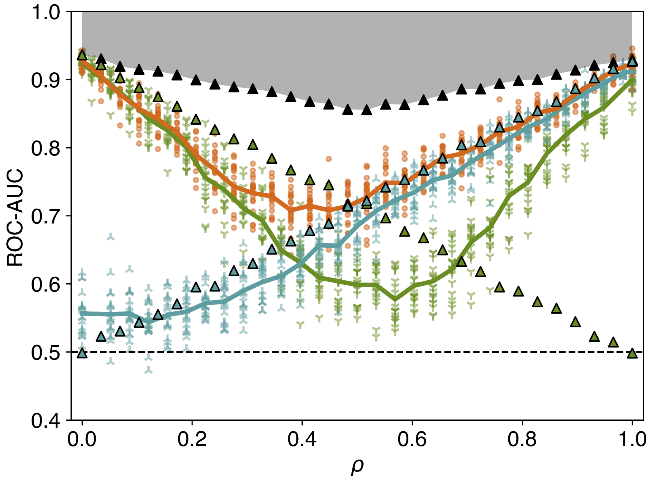


FIG. S14. Performance of missing link prediction on synthetic networks mixing SBM and assortative RGG networks with varying $\rho$ as shown in main text Fig. 3A, with partial theoretical ROC-AUC maximum curves shown for the RGG model (in blue) and the SBM model (in green).

### V. FOOD WEB DATA PROCESSING

The GlobAL daTabasE of traits and food Web Architecture (GATEWAy; [3, 4]; version 3, accessible at `doi.org/10.25829/idiv.283-3-756`) includes 222,151 consumer-resource interactions across 290 food web datasets spanning terrestrial above and below-ground, lake, marine, and stream ecosystem types (Figure S15A, Table S7). The consumer and resource nodes involved in each interaction are identified by the `con.taxonomy` and `res.taxonomy` columns respectively, and each interaction is associated with information for both the consumer and resource species on their respective lifestage, taxonomic aggregation level, movement type, metabolic type, and mass.

In constructing food webs from this list of interactions, we chose to additionally disaggregate food web nodes by lifestage to ensure the accuracy of trait values assigned to each node, e.g., we treated the adult and larval stages of the same species as separate nodes. Thus, unique nodes were identified by a combination of the `con.taxonomy` or `res.taxonomy` columns and the respective `lifestage` column. Lifestage disaggregation changed the set of nodes for 25 out of the 290 food webs (491 nodes added in total, 472 unique). We found nearly identical missing link prediction performance with or without lifestage disaggregationm for these food webs (Fig. S16).

In food web construction, we included all interactions labeled as *predacious*, *herbivorous*, *fungivorous*, *bacterivorous* and *detritivorous* (Fig. S15B). We excluded links labeled as *parasitic* or *parasitoid*, which applied to 2853 interactions in 20 food webs. In this database, the food webs containing parasitic interactions were mostly terrestrial aboveground food webs (Fig. S15C). We made this choice because parasites have been documented to change food web structure [6, 12, 21, 25, 34], and our ecologically-oriented topological predictors were not designed with parasitic interaction patterns in mind. Parasitic interactions are also not often systematically included in food webs, and if they are, their interactions are often inferred rather than directly observed [35]. We additionally excluded 1920 interactions with *NA* interaction type in the Carpinteria food web [24] as, upon inspection, these interactions involved parasites, though we included the interactions labeled as *NA* interaction type in the food webs from Refs. [23] and [40], as these links appeared to be non-parasitic trophic interactions. We included all interaction classifications: individual-based intact and attack (*ibi*), interactions with groups such as filter feeding or grazing (*nibi*), and *NA*. We combined the possible values for node taxonomic level into 4 categories: *species* ('species', 'subspecies', 'variety', 'form'), *genus* ('genus'), *family+* ( 'family', 'order', 'class', 'phylum', 'kingdom'), and *unclassified* ('NA', 'unranked'). There was only one case of a taxonomic level conflict between repeated records for the same unique node (node name *Gomphonema*, lifestage *NA*, taxonomic level records as *species* and *genus*), which we manually set to *genus* (Figure S15D).

We additionally excluded cannibalistic interactions (self-loops), which applied to 3275 interactions (1 of which was also parasitic) across 271 food webs, because we set up our missing link prediction approach for networks without self-loops, although we note this as an area for future work. We also trimmed any remaining multi-edges, or interactions recorded twice for the same two nodes in a food web, which applied to 307 interactions across 16 food webs. After trimming interactions, the dataset included 213,797 interactions among 19077 nodes (5562 unique node names).

For each node, we included three species traits as node attributes: metabolic type, movement type, and log mass. We filled in values for 12 nodes with metabolic type *NA* as listed in Table S8, leaving 9 possible values for metabolic type across the database (Fig. S17A). We manually replaced movement type values for non-living nodes (node names: *benthic detritus_NA*, *carrion-detritus_NA*, *CPOM_NA*, *Detritus_adults*, *Detritus_NA*, *FPOM_NA*, *Marine detritus_NA*, *POM_NA*, *POM (detritus)_NA*, *terrestrial detritus_NA*, *Sediment_NA*, *Unidentified detritus_NA*, *Phytodetritus_NA*, *dead organic matter_NA*, *detritus_NA*, *leaf litter_NA*) with a new *other_nonliving* category, and otherwise did not investigate between-web differences in movement type values as we were more concerned with in-web consistency due to our approach of predicting missing links in individual food webs. We filled in values for 42 living nodes with movement type *NA* as listed in Supplemental Data File 1, leaving 7 possible values for movement type in the database (Fig. S17B). Most food webs had 2-5 distinct metabolic types present (Fig. S17C) and 3 or 4 distinct movement types present (Fig. S17D).

We included all interaction records for a given food web, including those that were trimmed, when collecting trait values for a node in that food web. Metabolic type and movement type traits were categorical and thus turned into binary attributes for each category, with only those types present in a given food web included as potential values for a node. If movement type records differed for a species within a food web we listed a '1' in all relevant columns for that node after removing *NA* values because there might be biological reasons for which a species differed in movement type between interactions (56 instances across 28 food webs).

If mass records for a node differed within a food web (18 nodes across 14 food webs) we saved as the mass value the mean across all mass records for that node in the food web, thus representing an expected value for that node in that food web. Mass values were also completely missing within a food web for 1371 nodes (279 unique names) across 143 food webs. For 579 of these nodes with missing mass and an 'unclassified' taxonomic level, we replaced their mass with the minimum mass observed in their food web. These nodes were generally exceptional in that they tended to represent groups of species where mass might more appropriately be treated as a *NA* value (for example, the nodes *roots*, *leaf litter*, or *algae*). However, using our link prediction approach, missing numeric attributes need to be imputed for all nodes before they are used as features in prediction and we did not want to lose topological information about feeding interactions in the food web by excluding these nodes. Thus, we chose to replace these with the minimum mass because we observed that these were generally primary producers or detritus nodes that we expected to be at the bottom of the food web, and thus we assigned them a low mass for predicting missing links following body mass ratio expectations for consumer-resource links [4]. We assumed trait-matching rules learned by the meta-level model would likely learn to point links from nodes of lower body mass to nodes of higher body mass. We excluded nodes with metabolic type *invertebrate* and taxonomic level *unclassified* from this replacement, instead using the approach below.

For the remaining 792 nodes missing mass, we filled in their mass values using guesses informed by prior knowledge, by averaging the mass values of similar nodes found in the database. This strategy was employed to represent how missing trait values might be realistically assigned to species in a food web dataset (e.g., as in Ref. [48]). Node mass values were replaced (i) first with the mean mass value of nodes with the same name (typically, those of the same species) seen in other food webs (110 replacements), then (ii) with the mean mass value of species with the same first part of the node name (splitting node names by spaces, thus typically these were nodes of the same genus seen in other food webs, 317 replacements), then (iii) with the mean value of nodes in the same food web with the same movement and metabolic type (75 replacements), then (iv) with the mean value of nodes in the same food web with the same metabolic type (188 replacements), then (v) with the mean value of nodes in the same food web with the same movement type (10 replacements), then (vi) with the mean value of nodes from other webs with the same metabolic type (89 replacements), and then finally (vii) with the mean value of nodes from other webs with the same movement type (3 replacements).

Because we used assumptions to fill in missing mass values, we further provided information to the predictive model by providing a binary indicator for each node indicating whether its mass was originally missing.

The distribution of final mass values across all nodes in all the food webs was extremely skewed with many low mass nodes—the mean value in the dataset across nodes was 7954.16g but the median was 0.0005g. The majority of species mass values were under 1g (17,127 nodes out of 19,077), with the smallest mass value in the dataset for the *bacteria* node ($6.8 \times 10^{-21}$g) in the Lough Hyne food web and the largest value (52,395,350g) for the *Balaenoptera acutorostrata* (common minke whale) node in the Kongsfjorden food web. Because of this characteristic skewed nature of body mass distributions driven by evolutionary processes [7], we thus included log (base 10) mass as a node attribute rather than raw mass values (Fig. S18) following prior work [4].

To contextualize predictive performance results across the database we saved features related to (i) metadata and data processing (12 features, Table S4), (ii) global network topology (5 features, Table S5), and (iii) network assortativity (7 features, Table S6) for each food web. The values for these 25 features across the 290 food webs is provided in Supplemental Data File 2 and the distributions across the database are visualized in Figures S19, S20, and S21.

The median percent of nodes originally missing mass in a food web was 0%, but this value was up to 64.8% (in the Sutton Stream food web) (Fig. S19A). Most food webs had many nodes that were resolved at higher than the species level — the median percent of nodes with a higher taxonomic level was 25%, though this value was down to 0% (Lake Malawi food web, Caribbean Reef food web) and up to 63.5% (Dutch Microfauna food web PlotB) (Fig. S19B). It was also common for food webs to have some nodes with an *unclassified* taxonomic level, with the median percent of nodes with an *unclassified* taxonomic level at 12.2% (ranging from 0% in the Tuesday Lake 1984, Tuesday Lake 1986, and Scottish Lake food webs to an outlier of 100% in the Lake Malawi food web) (Figure S19C). Most food webs did not have any parasitic interactions removed (median 0%), with the outlier of the Carpinteria food web at 69.4% (Fig. S19D). Most food webs had a handful of cannibalistic interactions removed (median 2.4%), with the largest value (26.9%) for the PC2P5 food web (Fig. S19E). The final number of attributes for nodes in a food web varied from 5 to 15, with most food webs having around 8 attributes per node (Fig. S19F) — this was equal to the number of unique movement types in the food web (binary indicators) plus the number of unique metabolic types in the food web (binary indicators), an additional binary indicator in the case of mass being originally missing for any nodes in the food web, and one numeric attribute for log mass. The distributions for minimum, maximum, mean, and median log mass values across the food webs reflected the observation that most node masses were very small (Fig. S19G-J). Universally positive mass skewness values (median 0.6722) also indicated that all food webs had right skewed mass distributions, with the least skewed distribution (0.1884) for the Lough Hyne food web and the most skewed distribution (1.6973) for the Bylot food web.

Global network topology features varied across the database. The median number of nodes was 43.5 (median log nodes 1.64), with the lowest number of nodes (10) in the SF1M2 food web and the highest number (606) in the Chesapeake Bay food web (Fig. S20A). The median average degree was 5.28, with the lowest average degree (1.15) in the AP1 food web and the highest average degree (32.63) in the Weddell Sea food web (Fig. S20B). The distribution of directed connectance values appeared approximately normal with a median connectance value of 0.1253, the minimum connectance (0.0231) in the Kongsfjorden food web and the maximum connectance (0.2975) in the Skipwith Pond food web (Fig. S20C). The clustering coefficients of undirected versions of the food webs varied dramatically, with a median clustering coefficient value of 0.2245, a minimum (0), indicating no undirected triangles in 12 food webs (Brook trout lake, Deep lake, Gull lake north, Indian lake, Owl Lake, Rock Lake, South Lake, Twin Lake East, Twin Lake West, Wolf Lake, Svalbard, and PP1l2), and a maximum (0.674) in the SEW18 food web (Fig. S20D). The distribution of modularity values appeared approximately normal with a median value of 0.217, a minimum value (0.015) in the Iceland stream IS7 August 2008 food web, and a maximum value (0.457) in the Kongsfjorden food web (Fig. S20E). As 0.3 is considered a modularity value above which there is significant community structure [8], this indicates that most food webs had a low to moderate level of community structure, with variation across the database.

Assortativity coefficients measure to what extent nodes with similar attributes or degree tend to interact more than those with differing attributes and are 1 for perfect assortative mixing (nodes only connect to those with the same trait), 0 for no assortative mixing, and -1 for perfect disassortative mixing (nodes only connect to those with different traits) [37]. There was variety across the dataset in terms of whether food webs were assortative or disassortative by node attribute and degree.

Assortativity values based on bin logged mass showed a bimodal distribution, with most food webs showing no to slightly positive assortativity based on mass and another group of food webs showing somewhat larger assortativity values, with the maximum value (0.223) for the HEW22 food web. Some food webs were also disassortative by mass, with the lowest value (-0.172) for the CGP1 food web (Fig. S21A). Food webs generally displayed no to weakly negative assortativity based on metabolic type (median -0.0001) and movement type (median -0.003) (Fig. S21B,C). There was a tail of food webs for movement type assortativity showing higher assortativity values, with the largest value (0.404) for the Dutch Microfauna food web Plot B. There were wide ranges of assortativity values for assortativities based on node degree. The median degree assortativity in to out was negative (-0.1612) as was the median degree assortativity out to in (-0.2706) (Fig. S21D,E). This indicates that generalist consumers tended to link to specialist resources and generalist resources tended to link to specialist consumers, however this differed across the dataset, for example the maximum assortativity value out to in was quite large (0.9289 in the Iceland stream IS8 August 2008 food web). Conversely, assortativity values were generally slightly positive for degree assortativity out to out (median 0.094) and in to in (median 0.1859) (Fig. S21F,G), indicating that generalist resources slightly tended to link with generalist resources and generalist consumers slightly tended to link to generalist consumers, although there were many disassortative examples as well.

Finally, to contextualize our results, we further investigated the distributions of each of these 25 features across the database partitioned into 5 sets of food webs based on ecosystem type. Notable patterns we observed included that aquatic food webs in the dataset were generally smaller (streams: median 1.93 log nodes; lakes: median 1.57 log nodes; marine: median 1.45 log nodes) than the terrestrial food webs in the dataset (terrestrial aboveground: median 2.09 log nodes; terrestrial belowground: median 2.09 log nodes). Terrestrial belowground food webs were more clustered (median clustering coefficient 0.582) than food webs of the other types, and they were also more assortative by mass (median mass assortativity of 0.1735) than food webs of other types. Marine food webs had more dramatically right-skewed mass distributions (median skewness 0.8763) than food webs of other ecosystem types. Details about how these distributions differed across ecosystem types are provided in Supplemental Data File 3.

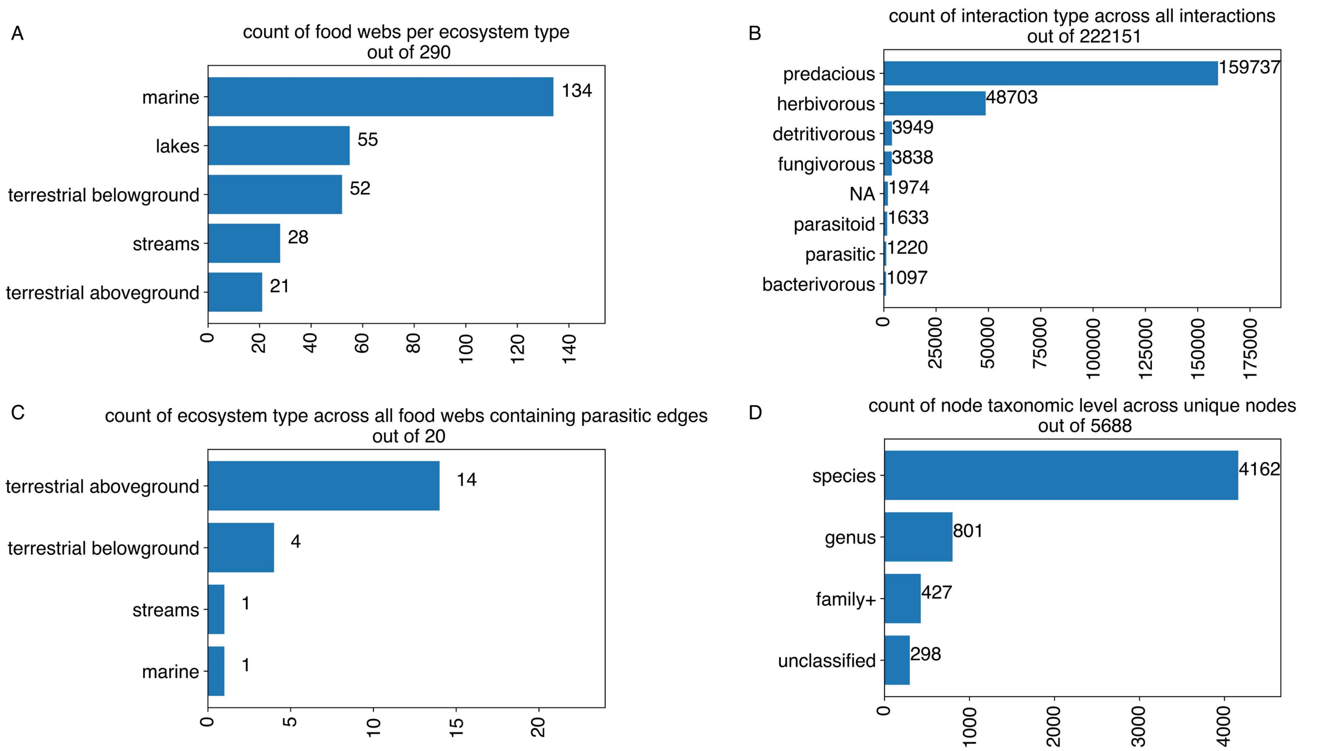


FIG. S15. Food web database details. (A) Ecosystem type counts across the 290 food webs. (B) Interaction type across the original 222,151 interactions. (C) Ecosystem type counts across the 20 food webs containing parasitic interactions. (D) Node taxonomic level counts across the 5688 unique node names in the original database.

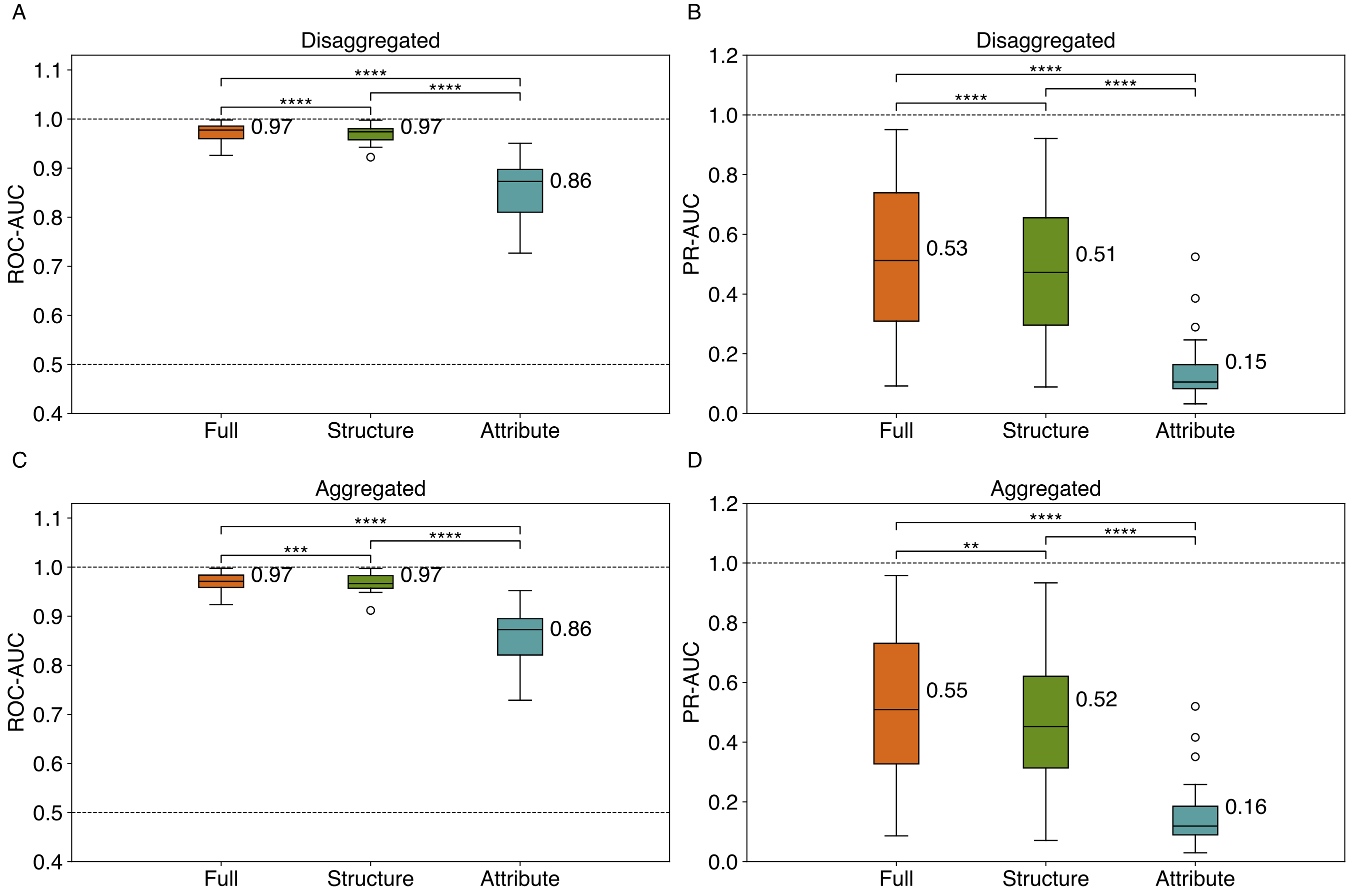


FIG. S16. Aggregated results for the 25 food webs whose node sets change with lifestage aggregation for stacked models using structure-only predictors ('structure'), attribute-only predictors ('attribute'), and both ('full'), (A) ROC-AUC and (B) PR-AUC results with nodes disaggregated by lifestage as in the main text results, (C) ROC-AUC and (D) PR-AUC results if nodes for the same species are aggregated across lifestages. Disaggregated results are averaged, as in the main text, over 5 iterations of evaluation across 5 unique folds (25 results per food web). Aggregated results are averaged across 5 unique folds (5 results per food web). Significant differences in mean model performance based on false discovery rate (FDR) adjusted (Benjamini-Hochberg method [2]) within-subjects pair-wise two-sided t-tests are shown where **** indicates a p-value<0.0001, *** p-value<0.001, and ** p-value<0.01.

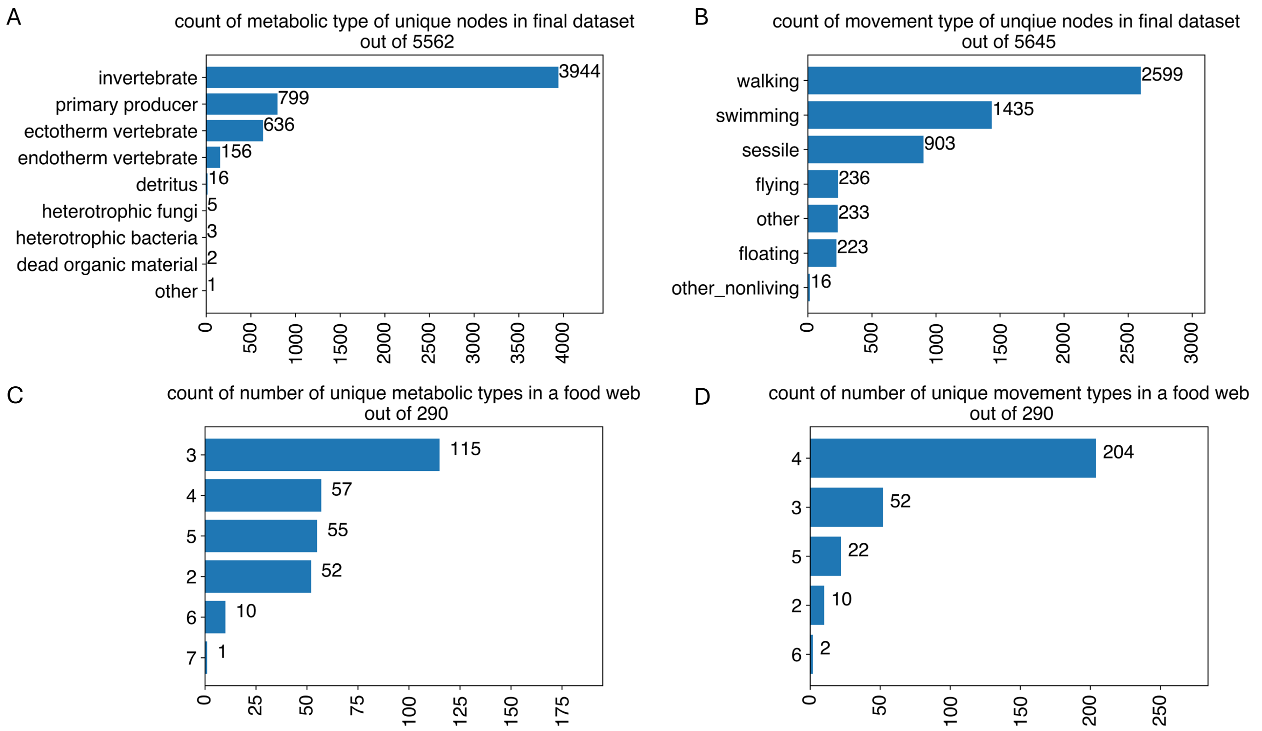


FIG. S17. Metabolic type and movement type details across the food web database.(A) Counts of node metabolic type across the 5562 unique nodes. (B) Counts of node movement type across the 5562 unique nodes, with multiple movement types either within or between webs for 83 nodes. (C) Counts of the number of final unique metabolic types in a given food web, across the 290 food webs. (D) Counts of the number of final unique movement types in a given food web, across the 290 food webs.

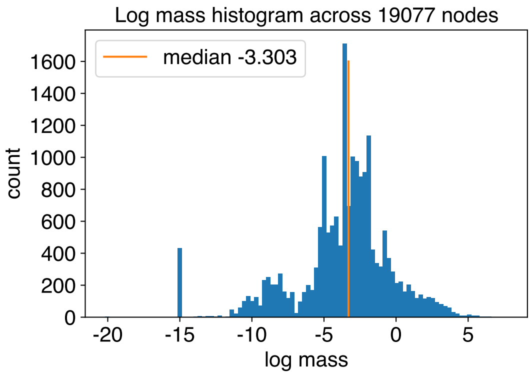


FIG. S18. Log mass histogram. Distribution of log (base 10) mass across the 19,077 nodes in the processed food web database.

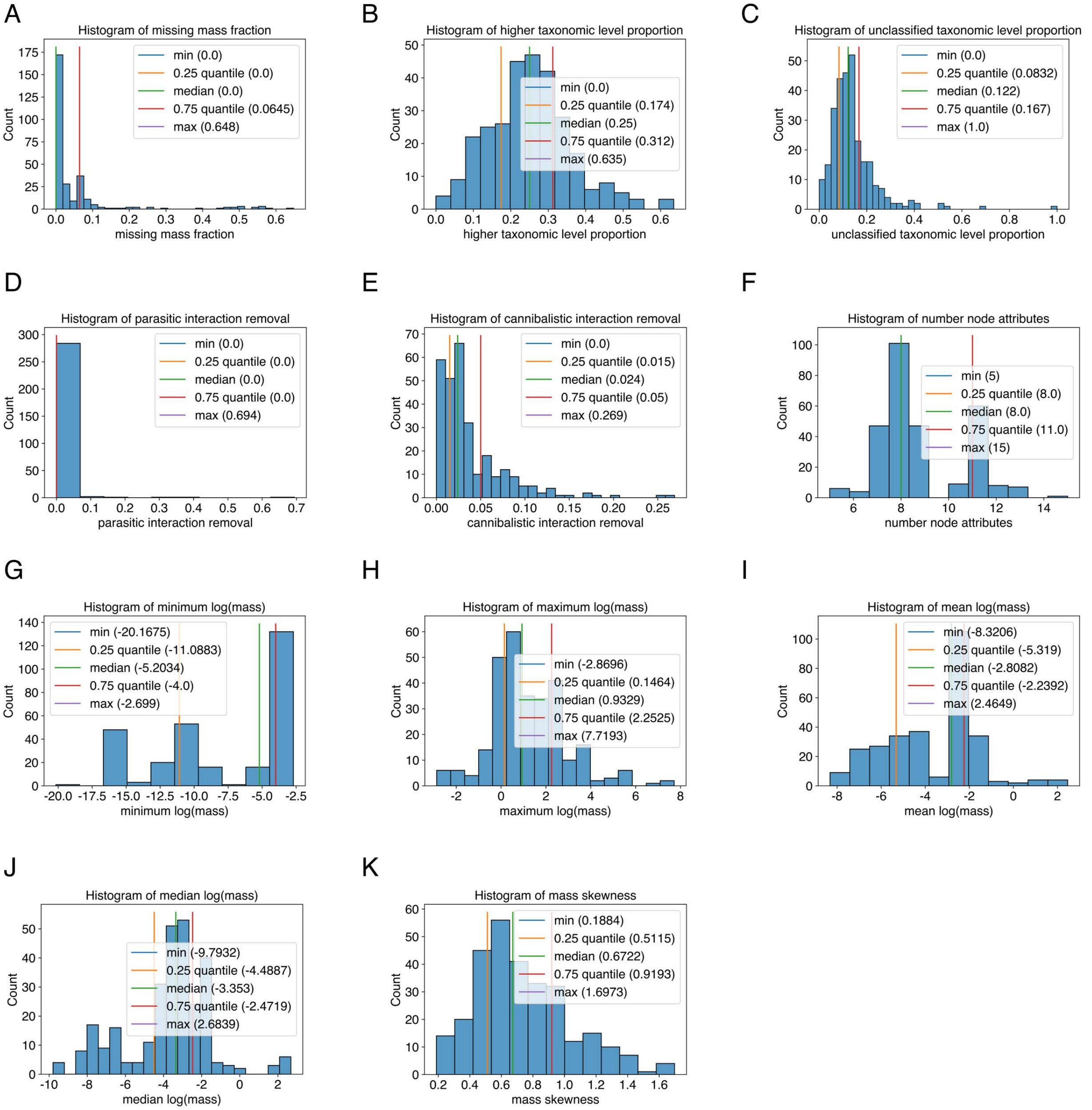


FIG. S19. Food web metadata and data processing feature distributions across the database. Histograms of (A) the missing mass fraction, (B) the higher taxonomic level proportion, (C) the unclassified taxonomic level proportion, (D) parasitic interaction removal, (E) cannibalistic interaction removal, (F) number node attributes, (G) minimum log mass, (H) maximum log mass, (I) mean log mass, (J) median log mass, and (K) mass skewness across the 290 food webs in the processed database.

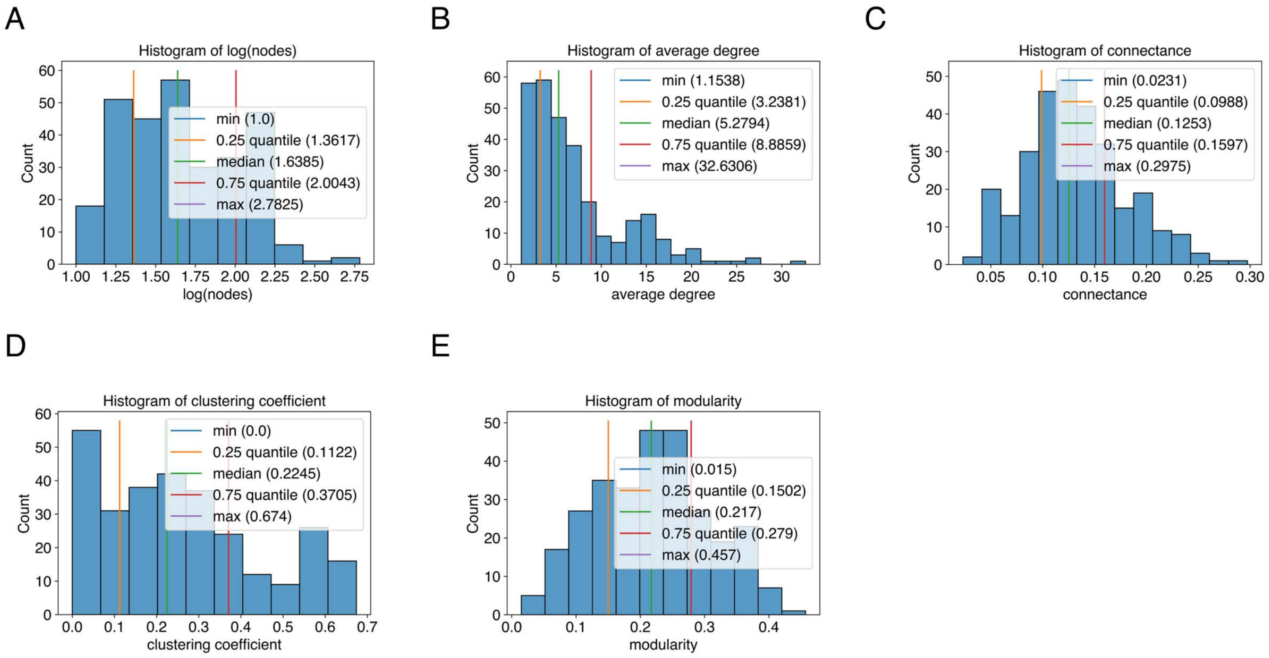


FIG. S20. Global network topology feature distributions across the database. Histograms of (A) log number of nodes, (B) average degree, (C) directed connectance, (D) clustering coefficient, and (E) modularity across the 290 food webs in the processed database.

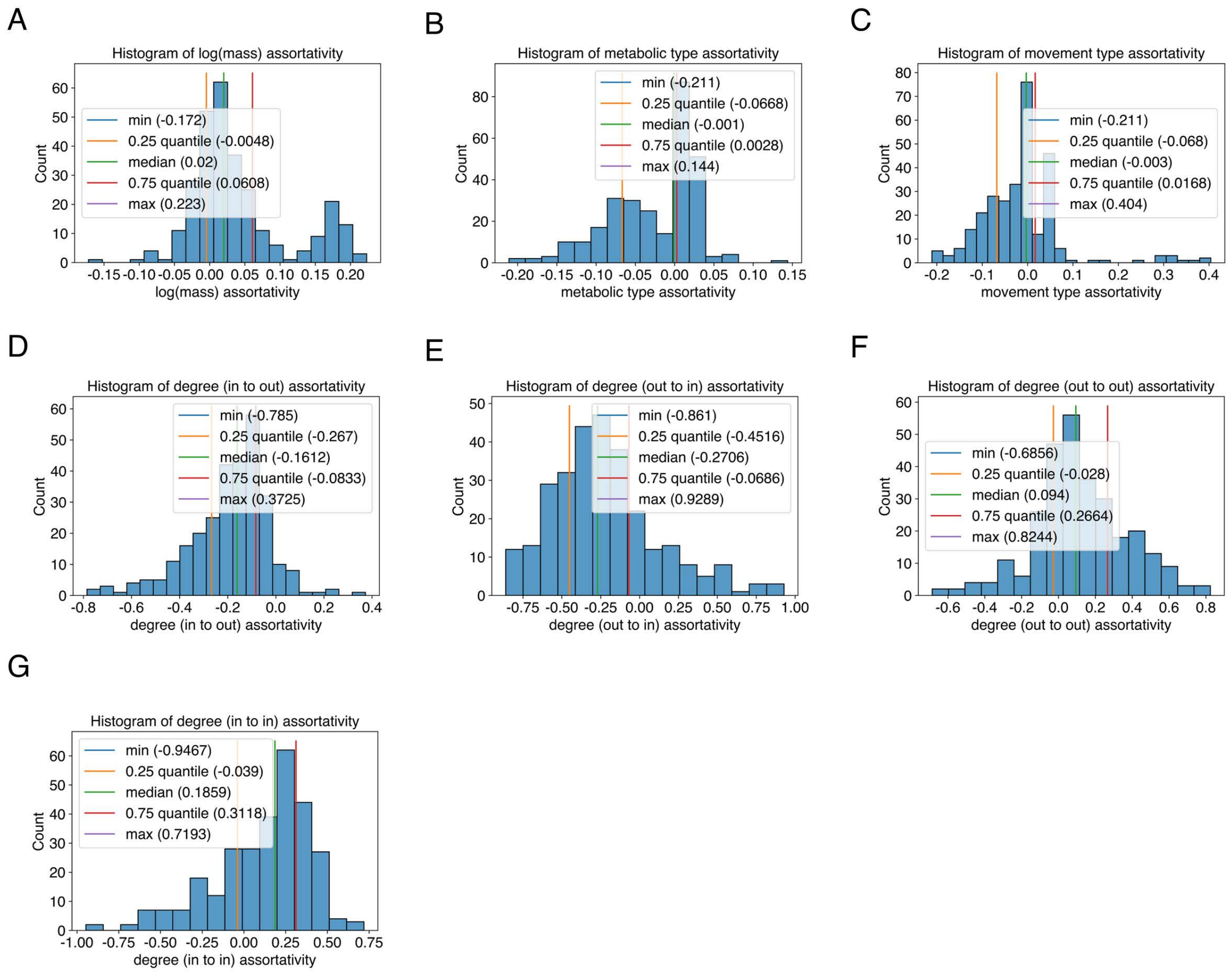


FIG. S21. Assortativity coefficient feature distributions across the database. Histograms of (A) log body mass assortativity, (B) metabolic type assortativity, (C) movement type assortativity, (D) degree (in to out) assortativity, (E) degree (out to in) assortativity, (F) degree (out to out) assortativity, and (G) degree (in to in) assortativity across the 290 food webs in the processed database.

### VI. SUPPLEMENTAL TABLES

TABLE S1: List of structure predictors. All of these predictors are included in the structure-only model and the full model.

| **Predictor Category** | **Name** | **Details** |
|---|---|---|
| Structure (pairwise) | Ecological common neighbor score | See Supplemental Equation (16), $\text{ECN}_{\text{score}}(i, j)$. |
| Structure (pairwise) | Ecological preferential attachment | See Main Text Equation (2), $\text{EPA}(i, j)$. |
| Structure (pairwise) | Personalized page rank | $j$th entry of the personalized page rank of node $i$, with input network treated as directed. Code: NetworkX [17]. |

| | | |
|---|---|---|
| Structure (node-level) | Page rank node $i$, Page rank node $j$ | Page rank values for nodes $i$ and $j$, with a directed network as input. Code: NetworkX [17]. |
| Structure (pairwise) | Ecological Jaccard coefficient | See Main Text Equation (10), EJC$(i, j)$. |
| Structure (pairwise) | Ecological Adamic/Adar index | See Supplemental Equation (10), EAA$(i, j)$. |
| Structure (pairwise) | Ecological resource allocation index | See Supplemental Equation (12), ERA$(i, j)$. |
| Structure (pairwise) | Shortest path | Shortest directed path between nodes $i$ and $j$. Ref: [30], Code: NetworkX [17]. |
| Structure (node-level) | Average neighbor in degree node $i$, Average neighbor in degree node $j$ | Code: NetworkX [17]. |
| Structure (node-level) | Average neighbor out degree node $i$, Average neighbor out degree node $j$ | Code: NetworkX [17]. |
| Structure (node-level) | In-degree centrality node $i$, In-degree centrality node $j$ | Code: NetworkX [17]. |
| Structure (node-level) | Out-degree centrality node $i$, Out-degree centrality node $j$ | Code: NetworkX [17]. |
| Structure (node-level) | Eigenvector centrality (in) node $i$, Eigenvector centrality (in) node $j$ | Eigenvector centralities in the in-direction for nodes $i$ and $j$. Code: NetworkX [17]. |
| Structure (node-level) | Eigenvector centrality (out) node $i$, Eigenvector centrality (out) node $j$ | Eigenvector centralities in the out-direction for nodes $i$ and $j$. Code: NetworkX [17]. |
| Structure (node-level) | Katz centrality (in) node $i$, Katz centrality (in) node $j$ | Katz centralities in the in-direction for nodes $i$ and $j$. Code: NetworkX [17]. |
| Structure (node-level) | Katz centrality (out) node $i$, Katz centrality (out) node $j$ | Katz centralities in the out-direction for nodes $i$ and $j$. Code: NetworkX [17]. |
| Structure (node-level) | Betweenness centrality node $i$, Betweenness centrality node $j$ | Shortest-path (directed) betweenness centralities for nodes $i$ and $j$. Code: NetworkX [17]. |
| Structure (node-level) | Closeness centrality (in) node $i$, Closeness centrality (in) node $j$ | Closeness centralities based on in-paths for nodes $i$ and $j$. Code: NetworkX [17]. |
| Structure (node-level) | Closeness centrality (out) node $i$, Closeness centrality (out) node $j$ | Closeness centralities based on out-paths for nodes $i$ and $j$. Code: NetworkX [17]. |
| Structure (node-level) | Load centrality node $i$, Load centrality node $j$ | Load centralities for nodes $i$ and $j$ based on shortest directed paths. Code: NetworkX [17]. |
| Structure (pairwise) | Ecological Leicht-Holme-Newman index | See Supplemental Equation (8), ELHN$(i, j)$. |
| Structure (pairwise) | Ecological common neighbor count | See Supplemental Equation (6), ECN$(i, j)$. |
| Structure (node-level) | Local clustering coefficient node $i$, Local clustering coefficient node $j$ | Local clustering coefficients based on directed triangles for nodes $i$ and $j$. Code: NetworkX [17]. |
| Structure (node-level) | Local triangles node $i$, Local triangles node $j$ | Local number of directed triangles for nodes $i$ and $j$. Code: NetworkX [17]. |
| Structure (pairwise) | Low rank approximation | Entry $i$, $j$ in low rank approximation (LRA) via singular value decomposition (SVD). Ref: [10], Code: [15]. |

| | | |
|---|---|---|
| Structure (pairwise) | Low rank dot product | Dot product of columns $i$ and $j$ in LRA via SVD for each pair of nodes $i$, $j$ (dLRA). Ref: [10], Code: [15]. |
| Structure (pairwise) | Low rank mean | Average of entries $i$ and $j$'s neighbors in LRA (mLRA). Ref: [10], Code: [15]. |
| Structure (pairwise) | Low rank approximation (approx) | An approximation of LRA. Ref: [10], Code: [15]. |
| Structure (pairwise) | Low rank dot product (approx) | An approximation of dLRA. Ref: [10], Code: [15]. |
| Structure (pairwise) | Low rank mean (approx) | An approximation of mLRA. Ref: [10], Code: [15]. |
| Structure (node-level) | Trophic level node $i$, Trophic level node $j$ | Ref: [29], Code: NetworkX [17]. |

TABLE S2: List of attribute predictors. All of these predictors are included in the attribute-only model and the full model.

| **Predictor Category** | **Name** | **Details** |
|---|---|---|
| Attribute (node-level) | $\{a\}$ node $i$, $\{a\}$ node $j$ | Raw attribute values (repeated for all $\|A\|$ attributes $a$). |
| Attribute (pairwise) | Euclidean distance | Euclidean distance between attribute vectors. |
| Attribute (pairwise) | Manhattan distance | Manhattan distance between attribute vectors. |
| Attribute (pairwise) | Cosine distance | Cosine distance between attribute vectors. |
| Attribute (pairwise) | Dot product | Dot product between attribute vectors. |
| Attribute (pairwise) | Euclidean (numeric) | Euclidean distance between numeric part of attribute vectors. |
| Attribute (pairwise) | Manhattan (numeric) | Manhattan distance between numeric part of attribute vectors. |
| Attribute (pairwise) | Cosine (numeric) | Cosine distance between numeric part of attribute vectors. |
| Attribute (pairwise) | Dot product (numeric) | Dot product between numeric part of attribute vectors. |
| Attribute (pairwise) | Hamming (binary) | Hamming distance (equivalent to Manhattan distance) between binary part of attribute vectors. |
| Attribute (pairwise) | Jaccard (binary) | Jaccard distance (equivalent to Tanimoto coefficient) between binary part of attribute vectors. |
| Attributes (pairwise) | $\{a\}$ ratio | $\frac{ai}{aj}$ repeated for all $\|N\|$ numeric attributes $a$. |

TABLE S3: List of nearest neighbor predictors. All of these predictors are the in full model and a subset based on $D5$ and $D6$ are in the structure-only model.

| **Predictor Category** | **Abbreviation** | **Definition** |
|---|---|---|
| Structure+Attribute (pairwise) | KNN (out) D1 | The fraction of out- neighbor sets of KNN_D1(i) in which $j$ is found, where D1 is the Euclidean distance between full min-max normalized node attribute vectors. |

| | | |
|---|---|---|
| Structure+Attribute (pairwise) | KNN (in) D1 | The fraction of in- neighbor sets of KNN_D1(j) in which $i$ is found, where D1 is the Euclidean distance between full min-max normalized node attribute vectors. |
| Structure+Attribute (pairwise) | KNN (out) D2 | The fraction of out- neighbor sets of KNN_D2(i) in which $j$ is found, where D2 is the Manhattan distance between full min-max normalized node attribute vectors. |
| Structure+Attribute (pairwise) | KNN (in) D2 | The fraction of in- neighbor sets of KNN_D2(j) in which $i$ is found, where D2 is the Manhattan distance between full min-max normalized node attribute vectors. |
| Structure+Attribute (pairwise) | KNN (out) D3 | The fraction of out- neighbor sets of KNN_D3(i) in which $j$ is found, where D3 is the Manhattan distance between the binary part of node attribute vectors. |
| Structure+Attribute (pairwise) | KNN (in) D3 | The fraction of in- neighbor sets of KNN_D3(j) in which $i$ is found, where D3 is the Manhattan distance between the binary part of node attribute vectors. |
| Structure+Attribute (pairwise) | KNN (out) D4 | The fraction of out- neighbor sets of KNN_D4(i) in which $j$ is found, where D4 is the Jaccard distance between the binary part of node attribute vectors. |
| Structure+Attribute (pairwise) | KNN (in) D4 | The fraction of in- neighbor sets of KNN_D4(j) in which $i$ is found, where D4 is the Jaccard distance between the binary part of node attribute vectors. |
| Structure (pairwise) | KNN (out) D5 | The fraction of out- neighbor sets of KNN_D5(i) in which $j$ is found, where D5 is the Jaccard similarity based on in-neighbor sets. |
| Structure (pairwise) | KNN (in) D5 | The fraction of in- neighbor sets of KNN_D5(j) in which $i$ is found, where D5 is the Jaccard similarity based on in-neighbor sets. |
| Structure (pairwise) | KNN (out) D6 | The fraction of out- neighbor sets of KNN_D6(i) in which $j$ is found, where D6 is the Jaccard similarity based on out-neighbor sets. |
| Structure (pairwise) | KNN (in) D6 | The fraction of in- neighbor sets of KNN_D6(j) in which $i$ is found, where D6 is the Jaccard similarity based on out-neighbor sets. |

TABLE S4. Food web metadata and data processing metrics used to understand predictive performance across the food web database.

| **Property** | **Type** | **Definition** |
|---|---|---|
| Ecosystem type | Categorical | Ecosystem type in the database. |
| Missing mass fraction | Numeric (range [0-1]) | Fraction of nodes in the network originally missing mass trait data. |
| Higher taxonomic level proportion | Numeric (range [0-1]) | Fraction of nodes in the network at a taxonomic level higher than species (‘genus’ or ‘family+’) |
| Unclassified taxonomic level proportion | Numeric (range [0-1]) | Fraction of nodes in the network with ‘unclassified’ taxonomic level (originally ‘NA’ or ‘unranked’) |
| Parasitic interaction removal | Numeric (range [0-1]) | The number of parasitic interactions removed divided by the final number of interactions in the network. |
| Cannibalistic interaction removal | Numeric (range [0-1]) | The number of cannibalistic interactions removed divided by the final number of interactions in the network. |
| Number node attributes | Numeric | The number of node attributes for this food web. |
| Minimum log mass | Numeric | The minimum log mass value in the food web. |
| Maximum log mass | Numeric | The maximum log mass value in the food web. |
| Mean log mas | Numeric | The mean log mass value across nodes in the food web. |
| Median log mass | Numeric | The median log mass value across nodes in the food web. |
| Mass skewness | Numeric | Pearsons 2nd coefficient of skewness of the raw mass values |

TABLE S5. Global network topology metrics used to understand predictive performance across the food web database. Topological properties were calculated using functions from the NetworkX library version 2.5.1 [17].

| **Property** | **Definition** |
|---|---|
| Log(nodes) | Where $n$ is the number of nodes, $log_{10}(n)$ |
| Average degree | Where $m$ is the number of directed edges, $m/n$ |
| Connectance | Number of directed edges divided by number of possible simple directed edges. Note that the denominator is adjusted such that cannibalistic interactions are not allowed. $m/(n*(n-1))$ |
| Clustering coefficient | The fraction of possible triangles present in the network. This calculation is performed on an undirected version of the network, thus ignoring directed structure of triangles. |
| Modularity | With communities found via greedy modularity maximization, this metric measures to what extent these communities have more connections within the same community as opposed to between communities. This calculation is performed on an undirected version of the network. [8] |

TABLE S6. Assortativity coefficients used to understand predictive performance across the food web database. Calculated using the NetworkX library version 2.5.1 [17], trait assortativity metrics are based on [37] and in- and out- degree assortativity coefficients are based on [14]. All assortativity coefficients take the directed structure of the network into account. In the rare cases of multiple movement types for a node in a network, one was randomly chosen.

| **Property** | **Definition** |
|---|---|
| Log body mass assortativity | Assortavity coefficient based on the log mass attribute split into 4 bins of equal size. |
| Metabolic type assortativity | Assortativity coefficient based on metabolic type categorical attribute. |
| Movement type assortativity | Assortativity coefficient based on movement type categorical attribute. |
| Degree (in to out) assortativity | Tendency of nodes with high in-degree (generalist consumers) to link to nodes with high out-degree (generalist resources). |
| Degree (out to in) assortativity | Tendency of nodes with high out-degree (generalist resources) to link to nodes with high in-degree (generalist consumers). |
| Degree (out to out) assortativity | Tendency of nodes with high out-degree (generalist resources) to link to nodes with high out-degree (generalist resources). |
| Degree (in to in) assortativity | Tendency of nodes with high in-degree (generalist consumers) to link to nodes with high in-degree (generalist consumers). |

TABLE S7: Food web details. Names, ecosystem types, and references for the 290 food webs in the GATEWAy database [3, 4]; (version 3, accessible at `doi.org/10.25829/idiv.283-3-756`).

| **Food web name** | **Ecosystem type** | **Ref** |
|---|---|---|
| Grand Caricaie marsh Clmown1 | terrestrial aboveground | [5] |
| Grand Caricaie marsh Clmown2 | terrestrial aboveground | [5] |
| Grand Caricaie marsh ClControl1 | terrestrial aboveground | [5] |
| Grand Caricaie marsh ClControl2 | terrestrial aboveground | [5] |
| Grand Caricaie marsh Scmown1 | terrestrial aboveground | [5] |
| Grand Caricaie marsh Scmown2 | terrestrial aboveground | [5] |
| Grand Caricaie marsh ScControl1 | terrestrial aboveground | [5] |
| Grand Caricaie marsh ScControl2 | terrestrial aboveground | [5] |
| Ythan Estuary | marine | [9] |
| AEW01 | terrestrial belowground | [11] |
| AEW02 | terrestrial belowground | [11] |
| AEW03 | terrestrial belowground | [11] |
| AEW04 | terrestrial belowground | [11] |
| AEW05 | terrestrial belowground | [11] |
| AEW06 | terrestrial belowground | [11] |
| AEW07 | terrestrial belowground | [11] |
| AEW08 | terrestrial belowground | [11] |
| AEW09 | terrestrial belowground | [11] |
| AEW11 | terrestrial belowground | [11] |
| AEW17 | terrestrial belowground | [11] |
| AEW18 | terrestrial belowground | [11] |
| AEW25 | terrestrial belowground | [11] |
| AEW27 | terrestrial belowground | [11] |
| AEW30 | terrestrial belowground | [11] |
| AEW49 | terrestrial belowground | [11] |
| HEW01 | terrestrial belowground | [11] |
| HEW02 | terrestrial belowground | [11] |
| HEW03 | terrestrial belowground | [11] |
| HEW04 | terrestrial belowground | [11] |
| HEW05 | terrestrial belowground | [11] |
| HEW06 | terrestrial belowground | [11] |
| HEW10 | terrestrial belowground | [11] |
| HEW11 | terrestrial belowground | [11] |

| | | |
|---|---|---|
| HEW12 | terrestrial belowground | [11] |
| HEW13 | terrestrial belowground | [11] |
| HEW16 | terrestrial belowground | [11] |
| HEW17 | terrestrial belowground | [11] |
| HEW21 | terrestrial belowground | [11] |
| HEW22 | terrestrial belowground | [11] |
| HEW36 | terrestrial belowground | [11] |
| HEW47 | terrestrial belowground | [11] |
| SEW01 | terrestrial belowground | [11] |
| SEW02 | terrestrial belowground | [11] |
| SEW03 | terrestrial belowground | [11] |
| SEW04 | terrestrial belowground | [11] |
| SEW05 | terrestrial belowground | [11] |
| SEW06 | terrestrial belowground | [11] |
| SEW07 | terrestrial belowground | [11] |
| SEW08 | terrestrial belowground | [11] |
| SEW09 | terrestrial belowground | [11] |
| SEW18 | terrestrial belowground | [11] |
| SEW35 | terrestrial belowground | [11] |
| SEW36 | terrestrial belowground | [11] |
| SEW37 | terrestrial belowground | [11] |
| SEW41 | terrestrial belowground | [11] |
| SEW43 | terrestrial belowground | [11] |
| SEW48 | terrestrial belowground | [11] |
| Kongsfjorden | marine | [13] |
| Alford lake | lakes | [18, 44] |
| Balsam lake | lakes | [18, 44] |
| Beaver lake | lakes | [18, 44] |
| Big hope lake | lakes | [18, 44] |
| Brandy lake | lakes | [18, 44] |
| Bridge brook lake | lakes | [18, 44] |
| Brook trout lake | lakes | [18, 44] |
| Buck pond | lakes | [18, 44] |
| Burntbridge lake | lakes | [18, 44] |
| Cascade lake | lakes | [18, 44] |
| Chub lake | lakes | [18, 44] |
| Chub pond | lakes | [18, 44] |
| Connera lake | lakes | [18, 44] |
| Constable lake | lakes | [18, 44] |
| Deep lake | lakes | [18, 44] |
| Emerald lake | lakes | [18, 44] |
| Falls lake | lakes | [18, 44] |
| Fawn lake | lakes | [18, 44] |
| Federation lake | lakes | [18, 44] |
| Goose lake | lakes | [18, 44] |
| Grass lake | lakes | [18, 44] |
| Gull lake | lakes | [18, 44] |
| Gull lake north | lakes | [18, 44] |
| Helldiver pond | lakes | [18, 44] |
| High pond | lakes | [18, 44] |
| Hoel lake | lakes | [18, 44] |
| Horseshoe lake | lakes | [18, 44] |
| Indian lake | lakes | [18, 44] |
| Little rainbow | lakes | [18, 44] |
| Long lake | lakes | [18, 44] |

| | | |
|---|---|---|
| Loon lake | lakes | [18, 44] |
| Lost lake | lakes | [18, 44] |
| Lost lake east | lakes | [18, 44] |
| Lower sister lake | lakes | [18, 44] |
| Oswego lake | lakes | [18, 44] |
| Owl lake | lakes | [18, 44] |
| Rat lake | lakes | [18, 44] |
| Razorback lake | lakes | [18, 44] |
| Rock lake | lakes | [18, 44] |
| Russian lake | lakes | [18, 44] |
| Safford lake | lakes | [18, 44] |
| Sand lake | lakes | [18, 44] |
| South lake | lakes | [18, 44] |
| Squaw lake | lakes | [18, 44] |
| Stink lake | lakes | [18, 44] |
| Twelfth tee lake | lakes | [18, 44] |
| Twin lake east | lakes | [18, 44] |
| Twin lake west | lakes | [18, 44] |
| Whipple lake | lakes | [18, 44] |
| Wolf lake | lakes | [18, 44] |
| Weddell Sea | marine | [20] |
| Tuesday Lake 1984 | lakes | [22] |
| Tuesday Lake 1986 | lakes | [22] |
| Chilean Intertidal Curaumilla | marine | [23] |
| Chilean Intertidal El Quisco | marine | [23] |
| Chilean Intertidal Las Cruces | marine | [23] |
| Chilean Intertidal Los Molles | marine | [23] |
| Chesapeake Bay | marine | (Kroll, unpublished) |
| Carpinteria | terrestrial aboveground | [24] |
| Scottish lake | lakes | [26] |
| Afon Hafren 2005 | streams | [26] |
| Allt a Mharcaidh | streams | [26] |
| Broadstone | streams | [26] |
| Dargall Lane | streams | [26] |
| Duddon Pike Beck | streams | [26] |
| Hardknott Gill | streams | [26] |
| Mill Stream | streams | [26] |
| Mosendale Beck | streams | [26] |
| Old Lodge | streams | [26] |
| Alert | terrestrial aboveground | [27] |
| Bylot | terrestrial aboveground | [27] |
| Herschel | terrestrial aboveground | [27] |
| Nenetsky | terrestrial aboveground | [27] |
| Svalbard | terrestrial aboveground | [27] |
| Yamal | terrestrial aboveground | [27] |
| Zackenberg | terrestrial aboveground | [27] |
| Gearagh | terrestrial aboveground | [32] |
| Dutch Detrital food web PlotA | terrestrial aboveground | [36, 42] |
| Dutch Detrital food web PlotB | terrestrial aboveground | [36, 42] |
| Dutch Detrital food web PlotC | terrestrial aboveground | [36, 42] |
| Bure Stream | streams | [16, 45] |
| Loddon Stream | streams | [16, 45] |
| Lyde Stream | streams | [16, 45] |
| Test Stream | streams | [16, 45] |
| Wensum Stream | streams | [16, 45] |

| | | |
|---|---|---|
| Lake Malawi | lakes | [38] |
| Iceland stream IS7 April 2009 | streams | [39] |
| Iceland stream IS7 August 2008 | streams | [39] |
| Iceland stream IS8 April 2008 | streams | [39] |
| Iceland stream IS8 August 2008 | streams | [39] |
| Caribbean Reef | marine | [40] |
| FloridaIslandE1 | terrestrial aboveground | [41, 43] |
| FloridaIslandE2 | terrestrial aboveground | [41, 43] |
| FloridaIslandE3 | terrestrial aboveground | [41, 43] |
| FloridaIslandE7 | terrestrial aboveground | [41, 43] |
| FloridaIslandE9 | terrestrial aboveground | [41, 43] |
| Lough Hyne | marine | [19] |
| Blackrock Stream | streams | [46] |
| Broad Stream | streams | [46] |
| Canton Creek | streams | [46] |
| Dempsters Stream | streams | [46] |
| German Creek | streams | [46] |
| Healy Creek | streams | [46] |
| Kye Burn | streams | [46] |
| Little Kye Burn | streams | [46] |
| Stony Stream | streams | [46] |
| Sutton Stream | streams | [46] |
| AP1 | marine | [33] |
| AP2 | marine | [33] |
| AP3 | marine | [33] |
| AP4 | marine | [33] |
| BP1 | marine | [33] |
| BP2 | marine | [33] |
| BP3 | marine | [33] |
| CGP1 | marine | [33] |
| CGP2 | marine | [33] |
| CGP3 | marine | [33] |
| CR1P1 | marine | [33] |
| CR1P2 | marine | [33] |
| CR1P3 | marine | [33] |
| CR1P4 | marine | [33] |
| CR2P1 | marine | [33] |
| CR2P2 | marine | [33] |
| CR2P3 | marine | [33] |
| CR2P4 | marine | [33] |
| F1P1 | marine | [33] |
| F1P2 | marine | [33] |
| F1P3 | marine | [33] |
| F1P4 | marine | [33] |
| F2P1 | marine | [33] |
| F2P2 | marine | [33] |
| F2P3 | marine | [33] |
| F2P4 | marine | [33] |
| FP1 | marine | [33] |
| FXAP1 | marine | [33] |
| FXAP2 | marine | [33] |
| FXAP3 | marine | [33] |
| FXAP4 | marine | [33] |
| FXBP1 | marine | [33] |
| FXBP2 | marine | [33] |

| | | |
|---|---|---|
| FXBP3 | marine | [33] |
| FXBP4 | marine | [33] |
| GJAP1 | marine | [33] |
| GJAP2 | marine | [33] |
| GJAP3 | marine | [33] |
| GJAP4 | marine | [33] |
| GJBP1 | marine | [33] |
| GJBP2 | marine | [33] |
| GJBP3 | marine | [33] |
| GJBP4 | marine | [33] |
| L1P1 | marine | [33] |
| L1P2 | marine | [33] |
| L1P3 | marine | [33] |
| L1P4 | marine | [33] |
| L2P1 | marine | [33] |
| L2P2 | marine | [33] |
| L2P3 | marine | [33] |
| L2P4 | marine | [33] |
| L3P1 | marine | [33] |
| L3P2 | marine | [33] |
| L3P3 | marine | [33] |
| L3P4 | marine | [33] |
| L4P1 | marine | [33] |
| L4P2 | marine | [33] |
| L4P3 | marine | [33] |
| L4P4 | marine | [33] |
| MBP1 | marine | [33] |
| MBP2 | marine | [33] |
| MBP3 | marine | [33] |
| MBP4 | marine | [33] |
| PC1P1 | marine | [33] |
| PC1P2 | marine | [33] |
| PC1P3 | marine | [33] |
| PC1P4 | marine | [33] |
| PC2P1 | marine | [33] |
| PC2P2 | marine | [33] |
| PC2P3 | marine | [33] |
| PC2P4 | marine | [33] |
| PC2P5 | marine | [33] |
| PGSBP1 | marine | [33] |
| PGSBP2 | marine | [33] |
| PGUBP1 | marine | [33] |
| PGUBP2 | marine | [33] |
| PGUBP3 | marine | [33] |
| PGUBP4 | marine | [33] |
| PP1I1 | marine | [33] |
| PP1I2 | marine | [33] |
| PP1I3 | marine | [33] |
| PP1I4 | marine | [33] |
| PP2I1 | marine | [33] |
| PP2I2 | marine | [33] |
| PP2I3 | marine | [33] |
| PP2I4 | marine | [33] |
| PP2M1 | marine | [33] |
| PP2M2 | marine | [33] |

| | | |
|---|---|---|
| PP2M3 | marine | [33] |
| PP2M4 | marine | [33] |
| RMP1 | marine | [33] |
| RMP2 | marine | [33] |
| RMP3 | marine | [33] |
| RMP4 | marine | [33] |
| RMP5 | marine | [33] |
| RV1P1 | marine | [33] |
| RV1P2 | marine | [33] |
| RV1P3 | marine | [33] |
| RV1P4 | marine | [33] |
| RV2P1 | marine | [33] |
| RV2P2 | marine | [33] |
| RV2P3 | marine | [33] |
| RV2P4 | marine | [33] |
| SF1I1 | marine | [33] |
| SF1I2 | marine | [33] |
| SF1I3 | marine | [33] |
| SF1I4 | marine | [33] |
| SF1M1 | marine | [33] |
| SF1M2 | marine | [33] |
| SF1M3 | marine | [33] |
| SF1M4 | marine | [33] |
| SF2I1 | marine | [33] |
| SF2I2 | marine | [33] |
| SF2I3 | marine | [33] |
| SF2I4 | marine | [33] |
| SF2M1 | marine | [33] |
| SF2M2 | marine | [33] |
| SF2M3 | marine | [33] |
| SF2M4 | marine | [33] |
| SP1 | marine | [33] |
| WP1 | marine | [33] |
| WP2 | marine | [33] |
| WP3 | marine | [33] |
| WP4 | marine | [33] |
| Skipwith Pond | lakes | [47] |

TABLE S8. Filling in NA metabolic type values in the food web database. Node names are written [taxonomy]_[lifestage]

| **Node name** | **Food web** | **Filled metabolic type value** |
|---|---|---|
| Trichoptera_larvae | Grand Caricaie marsh dominated by Cladietum marisci, mown Clmown1 | invertebrate |
| Trichoptera_larvae | Grand Caricaie marsh dominated by Cladietum marisci, mown Clmown2 | invertebrate |
| plankton_NA | Chilean Intertidal Curaumilla | primary producer |
| plankton_NA | Chilean Intertidal El Quisco | primary producer |
| plankton_NA | Chilean Intertidal Las Cruces | primary producer |
| plankton_NA | Chilean Intertidal Los Molles | primary producer |
| Carcinus sp._adults | Chesapeake Bay | invertebrate |
| Portunus spinimanus_adults | Chesapeake Bay | invertebrate |
| Leiostomus xanthurus_juveniles | Chesapeake Bay | ectotherm vertebrate |
| Plecoptera_larvae | Chesapeake Bay | invertebrate |
| Trichoptera_larvae | Chesapeake Bay | invertebrate |
| detritus_NA | Skipwith Pond | detritus |

[1] Lada A Adamic and Eytan Adar. Friends and neighbors on the Web. *Social Networks*, 25(3):211–230, July 2003.

[2] Yoav Benjamini and Yosef Hochberg. Controlling the false discovery rate: a practical and powerful approach to multiple testing. *Journal of the Royal statistical society: series B (Methodological)*, 57(1):289–300, 1995.

[3] Ulrich Brose. GlobAL daTabasE of traits and food Web Architecture (GATEWAy) version 1.0, 2018.

[4] Ulrich Brose, Phillippe Archambault, Andrew D. Barnes, Louis-Felix Bersier, Thomas Boy, João Canning-Clode, Erminia Conti, Marta Dias, Christoph Digel, Awantha Dissanayake, Augusto A. V. Flores, Katarina Fussmann, Benoit Gauzens, Clare Gray, Johanna Häussler, Myriam R. Hirt, Ute Jacob, Malte Jochum, Sonia Kéfi, Orla McLaughlin, Muriel M. MacPherson, Ellen Latz, Katrin Layer-Dobra, Pierre Legagneux, Yuanheng Li, Carolina Madeira, Neo D. Martinez, Vanessa Mendonça, Christian Mulder, Sergio A. Navarrete, Eoin J. O'Gorman, David Ott, José Paula, Daniel Perkins, Denise Piechnik, Ivan Pokrovsky, David Raffaelli, Björn C. Rall, Benjamin Rosenbaum, Remo Ryser, Ana Silva, Esra H. Sohlström, Natalia Sokolova, Murray S. A. Thompson, Ross M. Thompson, Fanny Vermandele, Catarina Vinagre, Shaopeng Wang, Jori M. Wefer, Richard J. Williams, Evie Wieters, Guy Woodward, and Alison C. Iles. Predator traits determine food-web architecture across ecosystems. *Nature Ecology & Evolution*, 3(6):919–927, June 2019.

[5] Marie-France Cattin Blandenier. *Food web ecology: models and application to conservation.* PhD Thesis, Verlag nicht ermittelbar, 2004.

[21] Abigail Z. Jacobs, Jennifer A. Dunne, Cristopher Moore, and Aaron Clauset. Untangling the roles of parasites in food webs with generative network models. Preprint, arxiv:1505.04741, 2015.

[22] Tomas Jonsson, Joel E Cohen, and Stephen R Carpenter. Food webs, body size, and species abundance in ecological community description. *Advances in ecological research*, 36(36):1–84, 2005.

[23] Sonia Kéfi, Eric L. Berlow, Evie A. Wieters, Lucas N. Joppa, Spencer A. Wood, Ulrich Brose, and Sergio A. Navarrete. Network structure beyond food webs: mapping non-trophic and trophic interactions on Chilean rocky shores. *Ecology*, 96(1):291–303, January 2015.

[24] Kevin D. Lafferty, Andrew P. Dobson, and Armand M. Kuris. Parasites dominate food web links. *Proceedings of the*

*National Academy of Sciences*, 103(30):11211–11216, July 2006.

[39] Eoin J. O'Gorman, Doris E. Pichler, Georgina Adams, Jonathan P. Benstead, Haley Cohen, Nicola Craig, Wyatt F. Cross, Benoît O.L. Demars, Nikolai Friberg, Gísli Már Gíslason, Rakel Gudmundsdóttir, Adrianna Hawczak, James M. Hood, Lawrence N. Hudson, Liselotte Johansson, Magnus P. Johansson, James R. Junker, Anssi Laurila, J. Russell Manson, Efpraxia Mavromati, Daniel Nelson, Jón S. Ólafsson, Daniel M. Perkins, Owen L. Petchey, Marco Plebani, Daniel C. Reuman, Björn C. Rall, Rebecca Stewart, Murray S.A. Thompson, and Guy Woodward. Impacts of Warming on the Structure and Functioning of Aquatic Communities. In *Advances in Ecological Research*, volume 47, pages 81–176. Elsevier, 2012.

[40] S. Opitz. Trophic interactions in caribbean coral reefs. Technical Report 43, ICLARM, 1996.

[41] Denise Piechnik, Sharon P. Lawler, and Neo D. Martinez. Food-web assembly during a classic biogeographic study species trophic breadth corresponds to colonization order. *Oikos*, 117:665–674, 2008.

[42] Valentina Sechi, Lijbert Brussaard, Ron G. M. De Goede, Michiel Rutgers, and Christian Mulder. Choice of Resolution by Functional Trait or Taxonomy Affects Allometric Scaling in Soil Food Webs. *The American Naturalist*, 185(1):142–149, January 2015.

[43] Daniel S. Simberloff and Edward O. Wilson. Experimental Zoogeography of Islands: The Colonization of Empty Islands. *Ecology*, 50(2):278–296, March 1969.

[44] J.W. Sutherland. Adirondack Biota Project. Technical report, 1989.
